# Cancer-Specific Alterations in Nuclear Matrix Proteins Determined by Multi-omics Analyses of Ductal Carcinoma *in Situ*

**DOI:** 10.1101/2024.02.13.580215

**Authors:** Ali Almutairy, Abdullah Alhamed, Stephen G. Grant, Miranda J. Sarachine Falso, Billy W. Day, Colton R. Simmons, Jean J. Latimer

**Affiliations:** Department of Pharmacology & Toxicology, College of Pharmacy, Qassim University, Buraydah, Saudi Arabia; Nova Southeastern University, Barry and Judy Silverman College of Pharmacy, Department of Pharmaceutical Sciences, Fort Lauderdale, FL; AutoNation Institute for Breast Cancer Research and Care, Fort Lauderdale, FL; King Saud University, College of Pharmacy, Pharmacology & Toxicology Department, Riyadh, Saudi Arabia; Nova Southeastern University, Dr. Kiran C. Patel College of Osteopathic Medicine, Department of Public Health, Fort Lauderdale, FL; University of Pittsburgh, Graduate School of Public Health, Department of Environmental & Occupational Health; University of Pittsburgh, School of Medicine, Department of Pharmacology & Chemical Biology, Pittsburgh, PA; University of Pittsburgh, School of Pharmacy, Department of Pharmaceutical Sciences, Pittsburgh, PA; University of Pittsburgh, School of Medicine, Department of Obstetrics and Gynecology, Pittsburgh, PA

## Abstract

Breast cancer (BC) is the most common cancer affecting women in the United States. Ductal carcinoma *in situ* (DCIS) is the earliest identifiable pre-invasive BC lesion. Estimates show that 14 to 50% of DCIS cases progress to invasive BC. Our objective was to identify nuclear matrix proteins (NMP) with specifically altered expression in DCIS and later stages of BC compared to non-diseased breast reduction mammoplasty and a contralateral breast explant using mass spectrometry and RNA sequencing to accurately identify aggressive DCIS. Sixty NMPs were significantly differentially expressed between the DCIS and non-diseased breast epithelium in an isogenic contralateral pair of patient-derived extended explants. Ten of the sixty showed significant mRNA expression level differences that matched the protein expression. These 10 proteins were similarly expressed in non-diseased breast reduction cells. Three NMPs (RPL7A, RPL11, RPL31) were significantly upregulated in DCIS and all other BC stages compared to the matching contralateral breast culture and an unrelated non-diseased breast reduction culture. RNA sequencing analyses showed that these three genes were upregulated increasingly with BC progression. Finally, we identified three NMPs (AHNAK, CDC37 and DNAJB1) that were significantly downregulated in DCIS and all other BC stages compared to the isogenically matched contralateral culture and the non-diseased breast reduction culture using both proteomics and RNA sequencing techniques.

## Introduction

Breast cancer (BC) was the most common cancer affecting women in the United States in 2023 (Siegel et al., 2023). The estimated number of new cases of invasive breast cancer in 2023 exceeded 297,800, with more than 43,000 deaths (Siegel et al., 2023). Mammographic screening and detection have been identified as factors contributing to the reduced mortality of BC and increased incidence of ductal carcinoma *in situ* (DCIS) (Duffy et al., 2020; Siegel et al., 2023). DCIS is the earliest identifiable pre-invasive breast cancerous lesion that may progress to invasive BC. The estimated number of new DCIS cases in the United States exceeded 55,000 cases in 2023, highlighting that DCIS constitutes approximately 20-25% of all diagnosed breast cancer cases (Siegel et al., 2023; van Seijen et al., 2019).

Estimates show that 14 to 50% of DCIS cases will breach the duct and progress to invasive breast cancer (Collins et al., 2005; Erbas et al., 2006; Maxwell et al., 2018; Sanders et al., 2015). The clinical behavior of DCIS lesions is not entirely understood. It is currently impossible to differentiate between aggressive and indolent DCIS cases at diagnosis. All cases are treated. Conventional management includes surgery with or without radiation and endocrine therapy, which can lead to complications (Doke et al., 2018; Hong et al., 2018).

One of the hallmark events in cancer progression is morphological alterations in cancerous cells such as irregular size and shape of the nucleus and an increase in the nucleus-to-cytoplasmic ratio. The nuclear matrix is a fibrous scaffolding system that participates in maintaining the spatial arrangement of the genome and nuclear components (Zink et al., 2004). Characteristic alterations in the protein composition of the nuclear matrix of tumor cells have proven to be useful to identify tumor markers. Specific nuclear matrix protein (NMP) changes are associated with cancerous cells relative to normal cells (Spencer et al., 2001). Changes in certain NMPs have been associated with prostate, colon, and breast cancers (Sjakste et al., 2004). Such markers may help to accurately identify invasive DCIS, improve early detection of invasive BC and prevent overtreatment complications. Paramount to this goal is to establish biomarkers of DCIS, especially invasive DCIS, with a focus on the nucleus of the tumor cells.

In this study, we used a novel tissue-engineering system to establish model systems representative of non-diseased breast as well as different molecular subtypes of all stages of normal breast and BC, including DCIS (Latimer, 2000, 2002). In addition, we examined NMP differences between these systems at both the protein and RNA levels. We specifically studied NMPs that were differentially expressed between a novel DCIS patient-derived extended explant with an abnormal karyotype and a similar culture from the isogenically matched contralateral, non-diseased breast from the same patient with a normal karyotype. We quantitatively analyzed the NMPs using a mass spectrometry (MS)-based relative quantitative methodology to identify biomarkers that were altered in both DCIS and invasive BC cell lines and absent in the non-diseased contralateral counterpart and an independent normal breast explant. Proteomic biomarker candidates meeting these specific criteria were then analyzed at the level of RNA sequencing to determine if the differential mRNA levels matched changes in protein levels for future molecular biomarker development.

## Materials and Methods

### Chemicals

Unless otherwise noted, all chemicals were from Sigma-Aldrich (St. Louis, MO).

### Cell culture

The DCIS and contralateral breast tissue (unaffected breast) were obtained from a 35-40 year old woman of Middle Eastern ancestry who underwent a double mastectomy. An additional non-diseased breast explant, JL-BRL-6 was used as normal control and obtained from a reduction mammoplasty surgery carried out at UPMC Magee-Womens Hospital (Visus et al. 2011). Human BC tissues that represent all BC stages were obtained with consent from women who were undergoing surgeries at UPMC Magee-Womens Hospital under IRB 0609002. (**EV Table 1**).

**Table 1.**
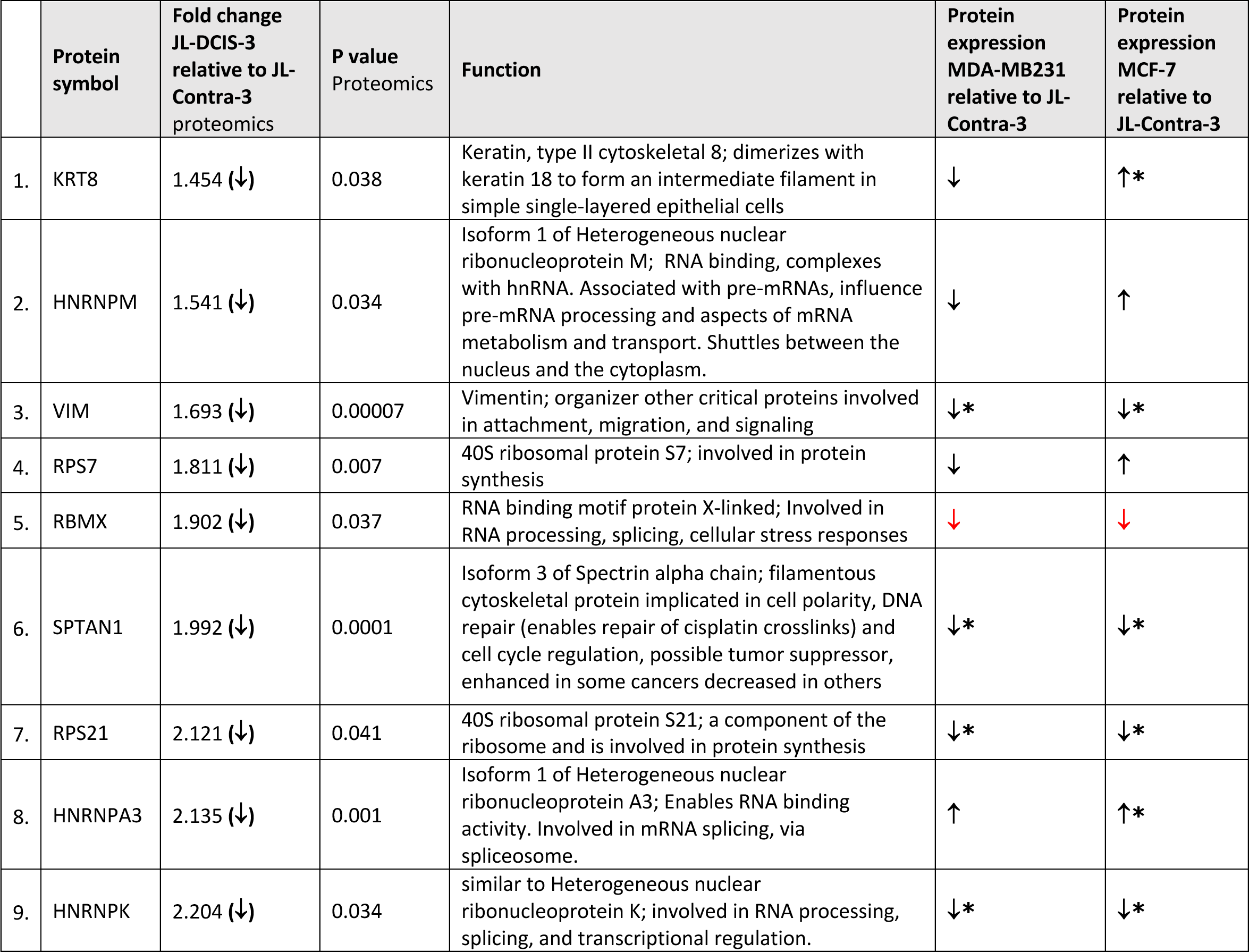
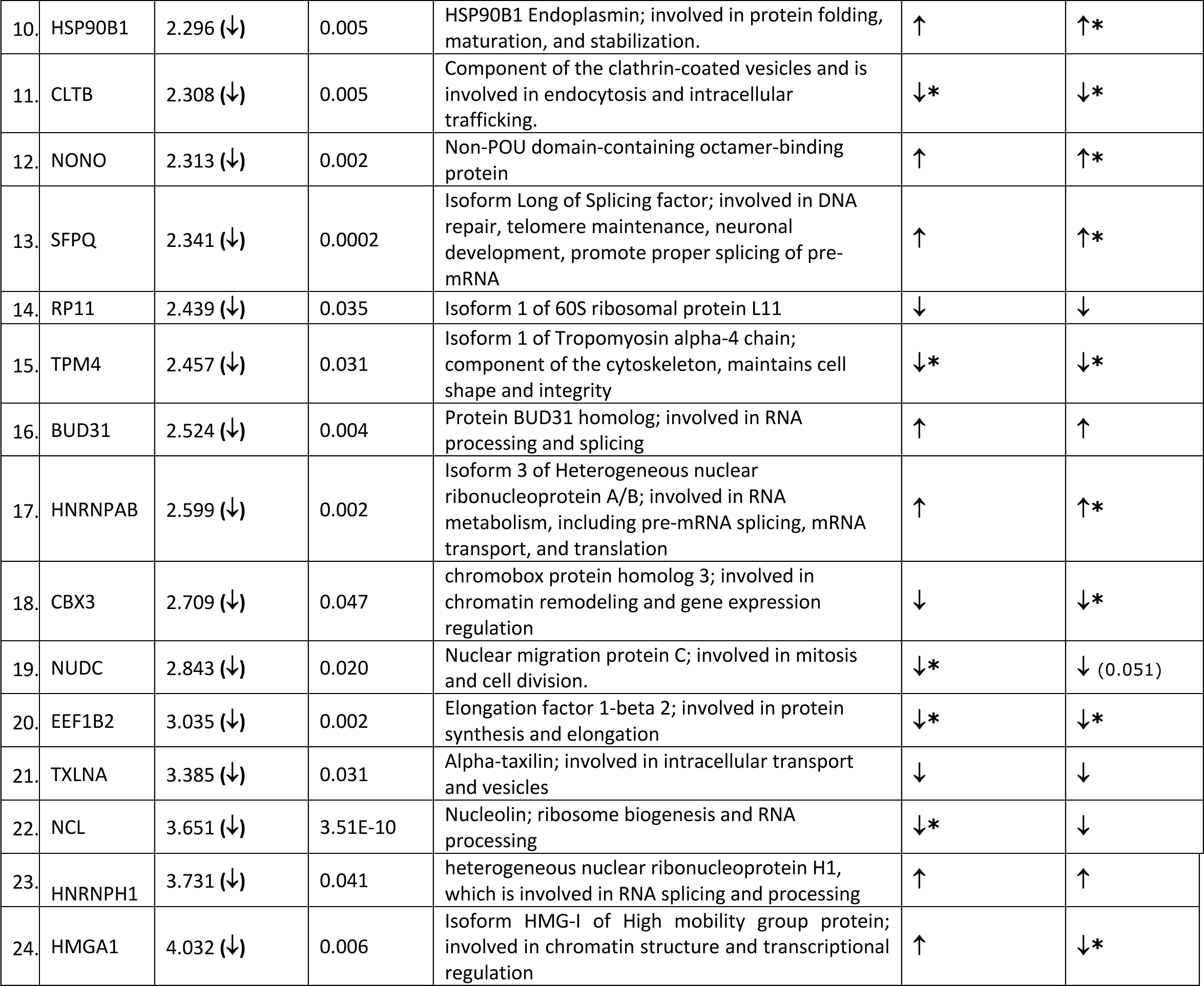
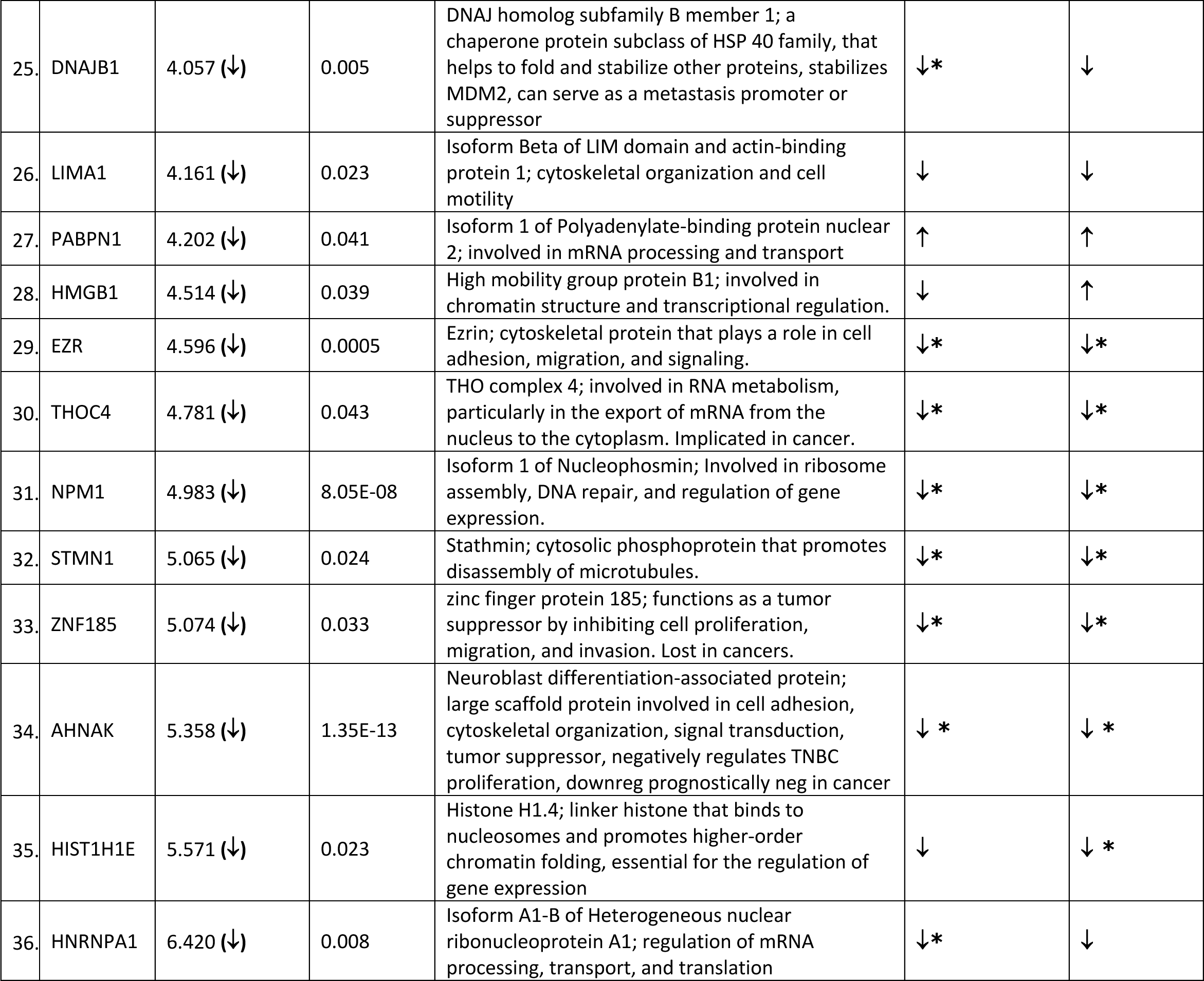
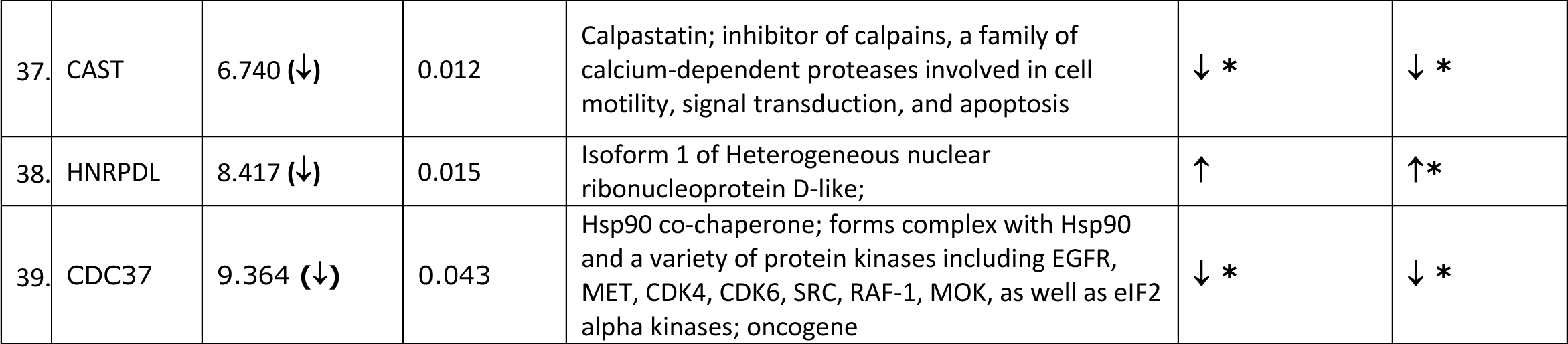
Significantly downregulated nuclear matrix proteins in JL-DCIS-3 relative to non-diseased JL-Contra-3 (all t tests 2-tailed).

|  | Protein symbol | Fold change JL-DCIS-3 relative to JL-Contra-3 proteomics | P value Proteomics | Function | Protein expression MDA-MB231 relative to JL-Contra-3 | Protein expression MCF-7 relative to JL-Contra-3 |
| --- | --- | --- | --- | --- | --- | --- |
| 1. | KRT8 | 1.454 (↓) | 0.038 | Keratin, type II cytoskeletal 8; dimerizes with keratin 18 to form an intermediate filament in simple single-layered epithelial cells | ↓ | ↑* |
| 2. | HNRNPM | 1.541 (↓) | 0.034 | Isoform 1 of Heterogeneous nuclear ribonucleoprotein M; RNA binding, complexes with hnRNA. Associated with pre-mRNAs, influence pre-mRNA processing and aspects of mRNA metabolism and transport. Shuttles between the nucleus and the cytoplasm. | ↓ | ↑ |
| 3. | VIM | 1.693 (↓) | 0.00007 | Vimentin; organizer other critical proteins involved in attachment, migration, and signaling | ↓* | ↓* |
| 4. | RPS7 | 1.811 (↓) | 0.007 | 40S ribosomal protein S7; involved in protein synthesis | ↓ | ↑ |
| 5. | RBMX | 1.902 (↓) | 0.037 | RNA binding motif protein X-linked; Involved in RNA processing, splicing, cellular stress responses | ↓ | ↓ |
| 6. | SPTAN1 | 1.992 (↓) | 0.0001 | Isoform 3 of Spectrin alpha chain; filamentous cytoskeletal protein implicated in cell polarity, DNA repair (enables repair of cisplatin crosslinks) and cell cycle regulation, possible tumor suppressor, enhanced in some cancers decreased in others | ↓* | ↓* |
| 7. | RPS21 | 2.121 (↓) | 0.041 | 40S ribosomal protein S21; a component of the ribosome and is involved in protein synthesis | ↓* | ↓* |
| 8. | HNRNPA3 | 2.135 (↓) | 0.001 | Isoform 1 of Heterogeneous nuclear ribonucleoprotein A3; Enables RNA binding activity. Involved in mRNA splicing, via spliceosome. | ↑ | ↑* |
| 9. | HNRNPK | 2.204 (↓) | 0.034 | similar to Heterogeneous nuclear ribonucleoprotein K; involved in RNA processing, splicing, and transcriptional regulation. | ↓* | ↓* |
| 10. | HSP90B1 | 2.296 (↓) | 0.005 | HSP90B1 Endoplasmin; involved in protein folding, maturation, and stabilization. | ↑ | ↑* |
| 11. | CLTB | 2.308 (↓) | 0.005 | Component of the clathrin-coated vesicles and is involved in endocytosis and intracellular trafficking. | ↓* | ↓* |
| 12. | NONO | 2.313 (↓) | 0.002 | Non-POU domain-containing octamer-binding protein | ↑ | ↑* |
| 13. | SFPQ | 2.341 (↓) | 0.0002 | Isoform Long of Splicing factor; involved in DNA repair, telomere maintenance, neuronal development, promote proper splicing of pre-mRNA | ↑ | ↑* |
| 14. | RP11 | 2.439 (↓) | 0.035 | Isoform 1 of 60S ribosomal protein L11 | ↓ | ↓ |
| 15. | TPM4 | 2.457 (↓) | 0.031 | Isoform 1 of Tropomyosin alpha-4 chain; component of the cytoskeleton, maintains cell shape and integrity | ↓* | ↓* |
| 16. | BUD31 | 2.524 (↓) | 0.004 | Protein BUD31 homolog; involved in RNA processing and splicing | ↑ | ↑ |
| 17. | HNRNPAB | 2.599 (↓) | 0.002 | Isoform 3 of Heterogeneous nuclear ribonucleoprotein A/B; involved in RNA metabolism, including pre-mRNA splicing, mRNA transport, and translation | ↑ | ↑* |
| 18. | CBX3 | 2.709 (↓) | 0.047 | chromobox protein homolog 3; involved in chromatin remodeling and gene expression regulation | ↓ | ↓* |
| 19. | NUDC | 2.843 (↓) | 0.020 | Nuclear migration protein C; involved in mitosis and cell division. | ↓* | ↓ (0.051) |
| 20. | EEF1B2 | 3.035 (↓) | 0.002 | Elongation factor 1-beta 2; involved in protein synthesis and elongation | ↓* | ↓* |
| 21. | TXLNA | 3.385 (↓) | 0.031 | Alpha-taxilin; involved in intracellular transport and vesicles | ↓ | ↓ |
| 22. | NCL | 3.651 (↓) | 3.51E-10 | Nucleolin; ribosome biogenesis and RNA processing | ↓* | ↓ |
| 23. | HNRNPH1 | 3.731 (↓) | 0.041 | heterogeneous nuclear ribonucleoprotein H1, which is involved in RNA splicing and processing | ↑ | ↑ |
| 24. | HMGA1 | 4.032 (↓) | 0.006 | Isoform HMG-I of High mobility group protein; involved in chromatin structure and transcriptional regulation | ↑ | ↓* |
| 25. | DNAJB1 | 4.057 (↓) | 0.005 | DNAJ homolog subfamily B member 1; a chaperone protein subclass of HSP 40 family, that helps to fold and stabilize other proteins, stabilizes MDM2, can serve as a metastasis promoter or suppressor | ↓* | ↓ |
| 26. | LIMA1 | 4.161 (↓) | 0.023 | Isoform Beta of LIM domain and actin-binding protein 1; cytoskeletal organization and cell motility | ↓ | ↓ |
| 27. | PABPN1 | 4.202 (↓) | 0.041 | Isoform 1 of Polyadenylate-binding protein nuclear 2; involved in mRNA processing and transport | ↑ | ↑ |
| 28. | HMGB1 | 4.514 (↓) | 0.039 | High mobility group protein B1; involved in chromatin structure and transcriptional regulation. | ↓ | ↑ |
| 29. | EZR | 4.596 (↓) | 0.0005 | Ezrin; cytoskeletal protein that plays a role in cell adhesion, migration, and signaling. | ↓* | ↓* |
| 30. | THOC4 | 4.781 (↓) | 0.043 | THO complex 4; involved in RNA metabolism, particularly in the export of mRNA from the nucleus to the cytoplasm. Implicated in cancer. | ↓* | ↓* |
| 31. | NPM1 | 4.983 (↓) | 8.05E-08 | Isoform 1 of Nucleophosmin; Involved in ribosome assembly, DNA repair, and regulation of gene expression. | ↓* | ↓* |
| 32. | STMN1 | 5.065 (↓) | 0.024 | Stathmin; cytosolic phosphoprotein that promotes disassembly of microtubules. | ↓* | ↓* |
| 33. | ZNF185 | 5.074 (↓) | 0.033 | zinc finger protein 185; functions as a tumor suppressor by inhibiting cell proliferation, migration, and invasion. Lost in cancers. | ↓* | ↓* |
| 34. | AHNAK | 5.358 (↓) | 1.35E-13 | Neuroblast differentiation-associated protein; large scaffold protein involved in cell adhesion, cytoskeletal organization, signal transduction, tumor suppressor, negatively regulates TNBC proliferation, downreg prognostically neg in cancer | ↓ * | ↓ * |
| 35. | HIST1H1E | 5.571 (↓) | 0.023 | Histone H1.4; linker histone that binds to nucleosomes and promotes higher-order chromatin folding, essential for the regulation of gene expression | ↓ | ↓ * |
| 36. | HNRNPA1 | 6.420 (↓) | 0.008 | Isoform A1-B of Heterogeneous nuclear ribonucleoprotein A1; regulation of mRNA processing, transport, and translation | ↓* | ↓ |
| 37. | CAST | 6.740 (↓) | 0.012 | Calpastatin; inhibitor of calpains, a family of calcium-dependent proteases involved in cell motility, signal transduction, and apoptosis | ↓ * | ↓ * |
| 38. | HNRPDL | 8.417 (↓) | 0.015 | Isoform 1 of Heterogeneous nuclear ribonucleoprotein D-like; | ↑ | ↑* |
| 39. | CDC37 | 9.364 (↓) | 0.043 | Hsp90 co-chaperone; forms complex with Hsp90 and a variety of protein kinases including EGFR, MET, CDK4, CDK6, SRC, RAF-1, MOK, as well as eIF2 alpha kinases; oncogene | ↓ * | ↓ * |

The explants were physically disaggregated into flasks containing a thin coat of Matrigel, a reconstituted basement membrane matrix (Biotechne, Minneapolis, MN). The explants were maintained in serum-rich MWRI, a previously described tissue culture medium (Latimer, 2000b, 2000a; Latimer et al., 2010). All the explants were incubated in a humidified atmosphere with 5% CO_2_ at 37 °C. Stages I-IV tumor samples were similarly processed and passaged to form patient-derived cultures (PDCs) (≤13 passages) and eventually to form PDC extended explants (PDCEEs) (>13 passages). PDCEEs were required of JL-DCIS-3, JL-Contra-3 and JL-BRL-6 because of the large number of cells required for the proteomics aspect of this study. Karyotypes are shown in **Fig EV1A-C**.

MDA-MB-231, a commercially available, stage IV, triple negative BC cell line, and MCF-7, a luminal type BC cell line, were both purchased from American Type Culture Collection (ATCC). Cells were cultured in DMEM supplemented with 10% heat-inactivated FBS and 1% penicillin-streptomycin.

Total RNA derived from PDCEE of stage I (JL-BTL-8, -4, -33), Stage II (JL-BTL-29, - 10) (Wend et al., 2013), Stage III (JL-BTL-12) (Sajithlal et al., 2010), and Stage IV (JL-BTL-21 and -60) were used for RNA expression comparisons.

### Nuclear Matrix Isolation

Forty 25-cm^2^ flasks of each cell line were required for nuclear matrix isolation and subsequent proteomics assessment. The nuclear matrix was isolated according to the method of Fey and Penman (Fey, 1988). Cells were incubated in 0.5% Triton X-100 in a buffered solution with 2 mM vanadyl ribonucleoside, an RNase inhibitor, for 10 min on ice to release lipids and soluble proteins. The remaining sample was pelleted at 1800 rpm at 4°C for 10 min and incubated in ammonium sulfate (0.25 M) with 2 mM vanadyl ribonucleoside for 10 min on ice. This step was performed as a salt extraction to release soluble cytoskeletal elements. The remaining sample was then pelleted at 1800 rpm at 4°C for 10 min. DNase I treatment was performed to remove soluble chromatin for 30 min at room temperature and the sample was pelleted at 2200 rpm at 4°C for 10 min. RNase A was added to remove RNA with a 10 min incubation at room temperature and then the sample was pelleted at 2200 rpm at 4°C for 10 min. Intermediate filaments and NMPs were then disassembled with 8 M urea and the insoluble carbohydrates and extracellular matrix components were pelleted with ultracentrifugation at 50,000 rpm for 1 h at 15°C. Dialysis was performed overnight in an assembly buffer containing KCl and imidazole-HCl to remove the urea and reassemble the intermediate filaments. Ultracentrifugation at 45,000 rpm for 90 min at 20°C was performed to pellet the intermediate filaments. The NMPs were precipitated with ethanol then quantified using the Coomassie (Bradford) Protein Assay (Thermo Scientific, Waltham, MA). All solutions contained 1 mM phenylmethylsulfonyl fluoride to inhibit serine proteases.

### iTRAQ Labeling

Protein analysis was performed by the method described by Ross et al. (2004). Briefly, 50 μg of precipitated NMPs from different samples were each resuspended in 20 μL of 0.5 M triethylammonium bicarbonate with 1 μL of 0.1% SDS. Tris(2-carboxyethyl) phosphine (TCEP, 5 mM, 2 μL) was then added to each sample and incubated at 60°C for 1 h. Thiols were then alkylated with 1 μL of 10 mM methyl methanethiosulfonate (MMTS) at room temperature for 10 min. Trypsin (10 μg) was then added to each sample and digestion was carried out at 37°C overnight. iTRAQ (isobaric tags for relative and absolute quantitation; AB Sciex, Framingham, MA) reagents were then added and incubated at room temperature for 2 h. After labeling, the samples were then pooled.

### OFFGEL Fractionation

The pooled iTRAQ sample was desalted using a C_18_ SepPak and then speed vacuumed to dryness. OFFGEL fractionation was performed. The 3100 OFFGEL Fractionator and the OFFGEL Kit pH 3-10 (Agilent Technologies, Santa Clara, CA) was used following a 24-well set up. An Immobiline DryStrip, pH 3-10, 24 cm (GE Healthcare Bio-Sciences, Piscataway, NJ) was used. Fifteen minutes before sample loading, the gel strip was rehydrated in the assembled device with 40 μL OFFGEL Rehydration solution per well. The iTRAQ peptides were resuspended in the Peptide OFFGEL solution to a final volume of 3.6 mL. The diluted sample (150 μL) was distributed into the 24 wells. The default OFFGEL peptide 24-cm strip program on the instrument was used with a maximum current of 50 μA until 50 kVh was reached. The fractions were then recovered from each well. Methanol (150 μL, 50% aqueous containing 0.1% trifluoroacetic acid [TFA]) was then added back to each well and they were left on the benchtop for 20 min. The solution was then recovered and added back to the appropriate fraction and the entire sample was speed vacuumed to dryness and resuspended in 0.1% aqueous TFA for nanoLC (Dionex, Sunnyvale, CA) separation.

### Nano-LC-MALDI-TOF/TOF-MS/MS

The OFFGEL-fractionated samples were further fractionated on an RP-LC Ultimate system (Dionex, Sunnyvale, CA). They were first loaded onto a trap column (300 μm i.d. x 5 mm, PepMap C18 100, 5 μm, 100 Å) and washed for 10 min with 2% acetonitrile (ACN), 0.1% TFA at a flow of 30 μL/min. They were then loaded onto an analytical column (75μm i.d. x 150 mm, Pep Map C18 100 material 3 μm, 100 Å) and fractionated using a gradient of 5-30% B in 110 minutes, 30-60% B in 60 minutes, and 60-100% B in 10 minutes with a flow rate of 250 nL/min. Solution A was 5% ACN, 0.1% TFA, and solution B was 85% ACN, 5% isopropanol (IPA), and 0.1% TFA. Five minutes after the sample injection, the Probot^TM^ Micro Fraction Collector (Dionex, Sunnyvale, CA) was signaled to start spotting. The Probot was used to collect 15-second spots on the ABI 4800 LC-MALDI metal target (AB Sciex, Framingham, MA) in a 16 x 48 array. A total of 768 spots were collected for each OFFGEL fraction, and two LC runs were done on each target. This resulted in a total of 12 plates. The μTee (Dionex, Sunnyvale, CA) mixer was used to co-spot the matrix α-cyano-4-hydroxycinnamic acid (CHCA; 7 mg) in 1 mL of 50% ACN, 0.1% TFA, with mM ammonium citrate and 10 fmol angiotensin II), delivered at a flow rate of 1.577 μL/min. For MALDI-TOF/TOF-MS/MS analysis (4800 Proteomics Analyzer; AB Sciex, Framingham, MA), MS spectra were acquired from 900 to 4000 Da with a focus mass of 2000 Da. MS processing was performed using the angiotensin II internal standard with a 250-ppm max outlier error. Up to 10 peaks were selected for peptide sequencing by MS/MS. Peptide CID (air) was performed at 2 kV.

### Protein Identification, Quantification and Statistical Analysis

The Paragon algorithm in ProteinPilot^TM^ Software 2.0 (AB Sciex, Framingham, MA) was used for protein identification. Proteins were identified by searching against the IPI database v 3.46. Searched results were processed with the Pro Group algorithm (AB Sciex, Framingham, MA). Search parameters included iTRAQ labeling of the N-terminus and lysine residues, cysteine modification by MMTS, and digestion by trypsin. Isoform-specific identification and quantification was done by excluding all shared peptides and including only unique peptides. The p-value calculated by the software was used to determine the significance of observed differences in protein expression. This p-value tests the null hypothesis that the actual protein ratio is 1:1 and that the observed protein ratio is different than that only by chance. Only proteins identified with >95% confidence or ProtScore > 1.3 were selected for analysis.

### RNA Extraction and Quantification

Two 25-cm^2^ flasks of each cell line were harvested for RNA extraction. Total RNA was extracted with Qiazol reagent using the miRNeasy kit (Qiagen, Germantown, MD) following the manufacturer’s protocol. The quality of total RNA was evaluated using an RNA ScreenTape system using a 4200 TapeStation instrument (Agilent, Santa Clara, CA; C2991AA). RNA-sequencing libraries were prepared using Illumina® TruSeq stranded Total RNA Library Prep Gold (Illumina, Foster City, CA) and were sequenced using Illumina® NextSeq 500 High Output v2 Kit to obtain 150-bp paired-end reads. The sequencing depth for each sample was approximately 45 million reads.

The RNA sequencing raw data were analyzed using Partek®Flow® Software, version 10.0 (Partek Inc., 2022). The reads were aligned into a *Homo sapiens* reference genome (hg38) using the annotation model Ensembl transcripts release 105 (European Bioinformatic Institute, Hinxton, U.K.). Transcript abundance was expressed in fragments per kb of transcript per million mapped reads (FPKM). Differential gene expression analyses were done with Microsoft Excel (version 16.63) and GraphPad Prism (version 9.3.1).

### RNAsequencing Analyses

RNA sequencing results were based on three independent RNA samples and were expressed as the mean ± standard error for each group. Two-tailed Student’s *t*-tests were performed using Microsoft Excel (version 16.63) and GraphPad Prism software (version 9.3.1). A p-value of < 0.05 was considered to indicate a statistically significant difference.

## Results

The study design was based on a comparison of DCIS (JL-DCIS-3), a patient-derived culture extended explant (PDCEE), and an isogenically matched non-diseased contralateral culture (JL-Contra-3). These explants represent rare *in vitro* model systems established by directly culturing human DCIS and contralateral tissue in a system developed using a thin coat of Matrigel and stem cell-based medium (Latimer, 2000b; Latimer et al., 2010). Karyotyping of JL-DCIS-3 showed it was 46XX but contained a derivative chromosome 14, whereas JL-Contra-1 had a normal 46XX karyotype (**Fig EV1A, B**).

Gene expression in a breast reduction mammoplasty PDCEE established using the same culture conditions (JL-BRL-6) was compared with JL-Contra-3 to establish a non-diseased gene expression pattern. JL-BRL-6 also manifested a normal karyotype (**Fig EV1C**). The widely used commercial tumor cell lines MCF-7 and MDA-MB231 were included for comparison as commonly used standards to represent invasive BC for proteomic assessment of NMPs.

### Proteomic Analysis of NMPs

Proteomic analysis was performed on a total of five samples (MCF-7, MDA-MB-231, JL-BRL-6, JL-Contra-3 and JL-DCIS-3). Using the iTRAQ method, a total of 270 proteins were identified from the nuclear matrix of these samples out of approximately 1300 proteins previously identified as being associated with the nucleus (Dellaire, 2003). Of the 270 NMP proteins, we identified 60 whose levels significantly differed between JL-DCIS-3 and JL-Contra-3 (**Tables 1 and 2**). Thirty-nine of these proteins were downregulated in JL-DCIS-3 relative to JL-Contra-3 (Table 1) and 21 of these proteins were upregulated in JL-DCIS-3 relative to JL-Contra-3 (**Table 2**).

**Table 2.**
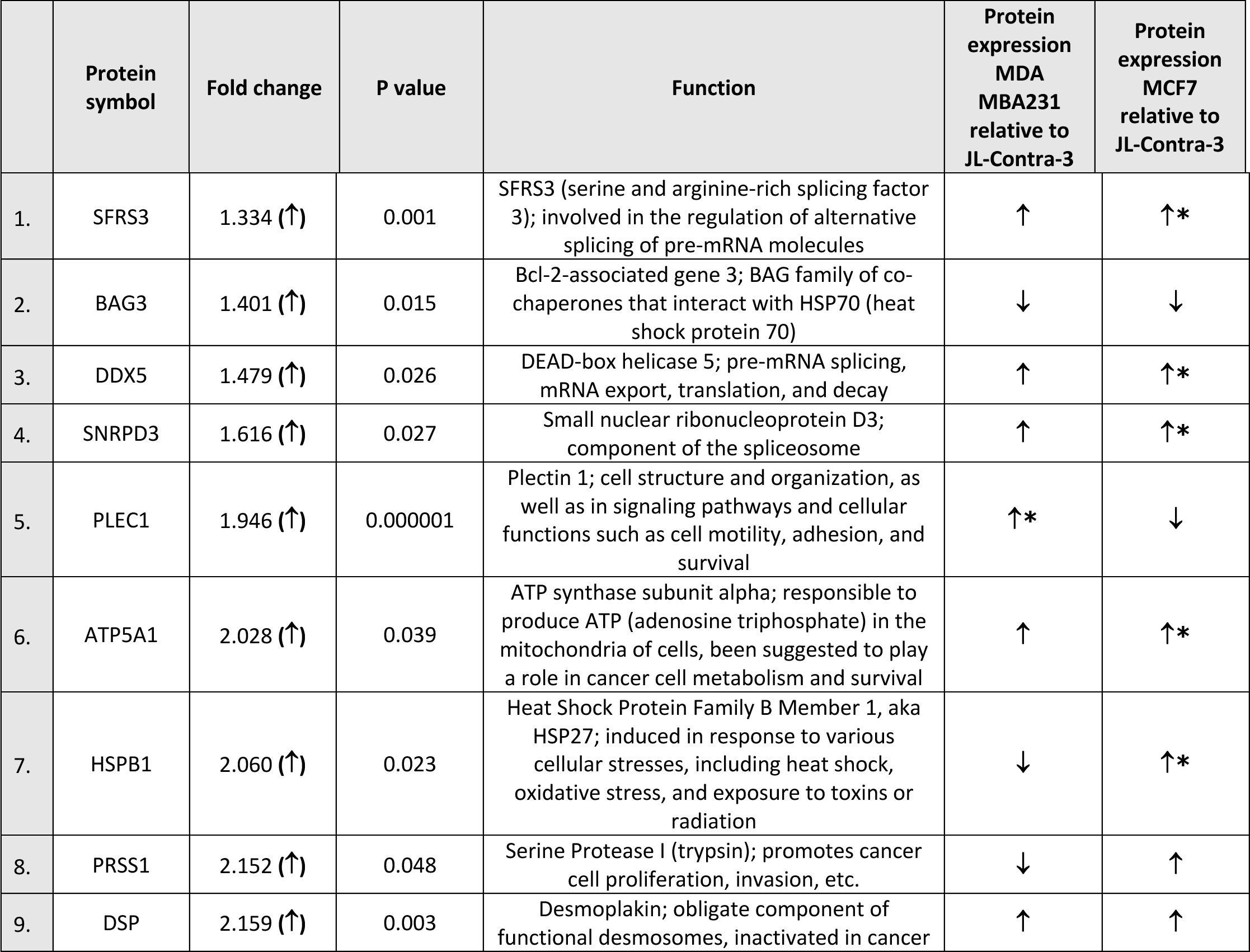
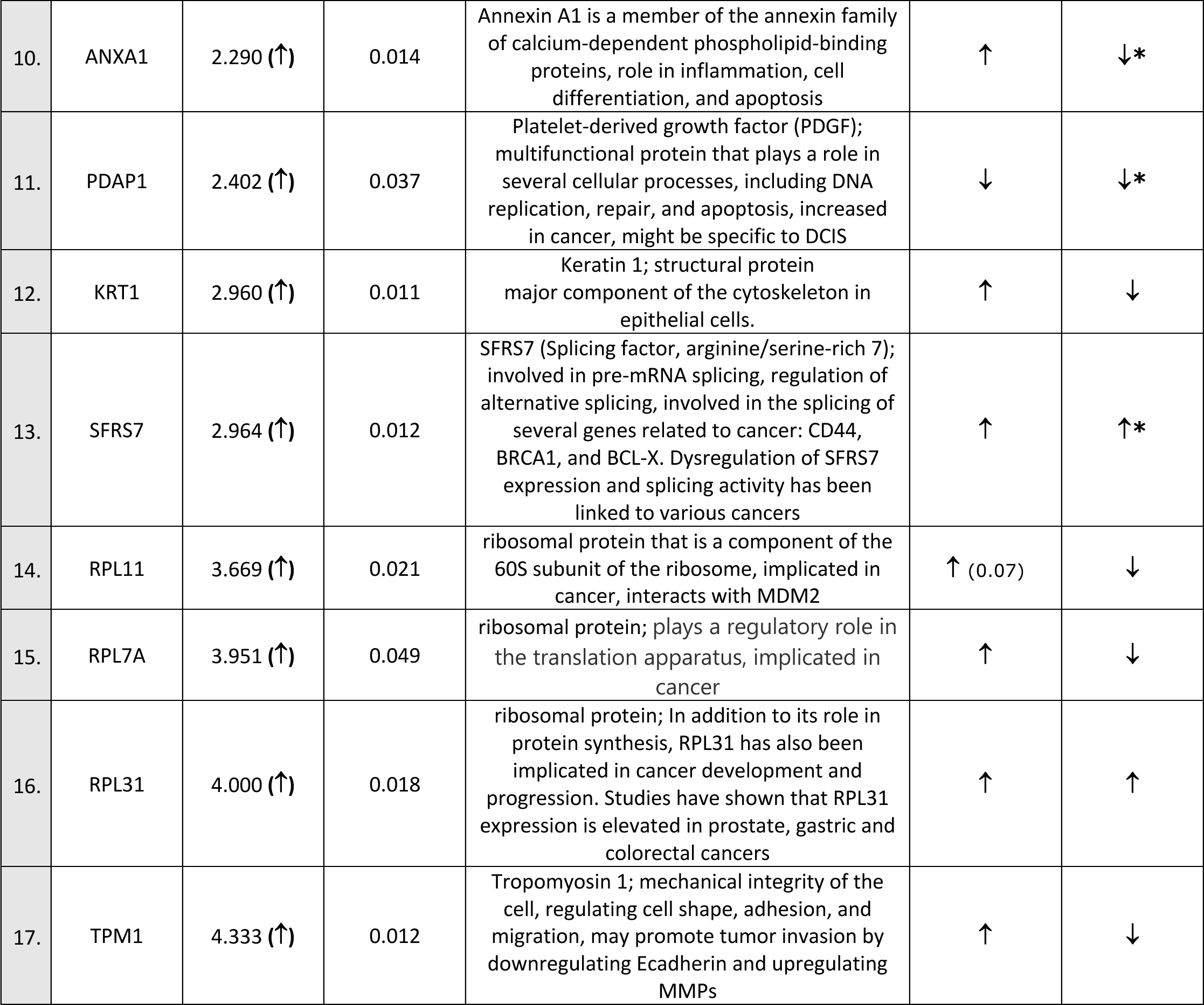
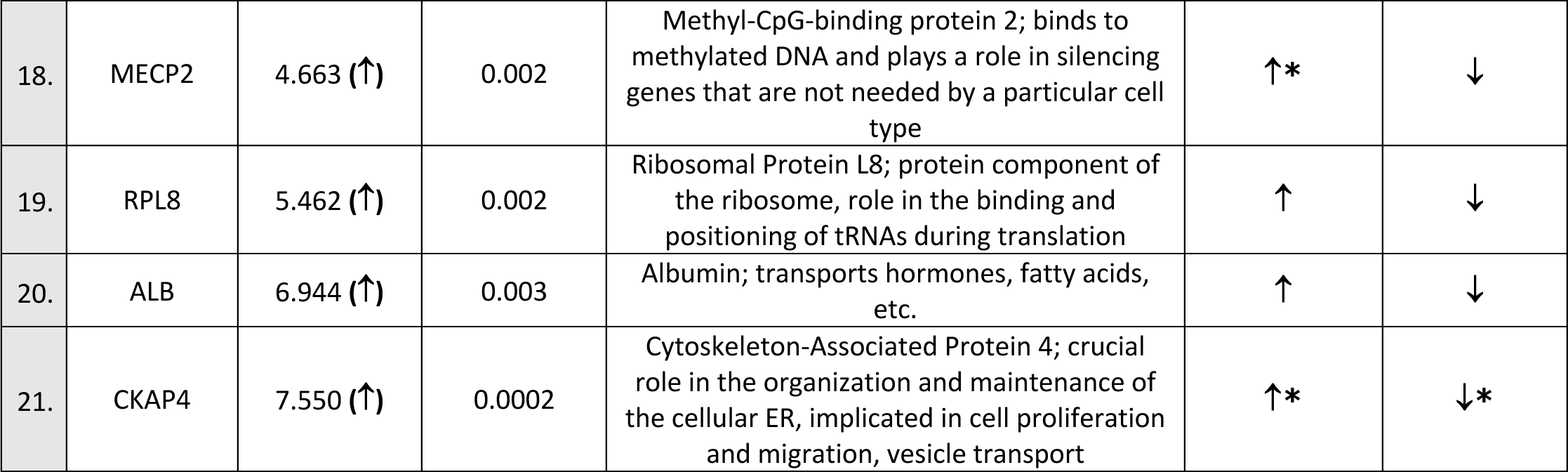
Significantly upregulated nuclear matrix proteins in JL-DCIS-3 relative to non-diseased JL-Contra-3.

|  | Protein symbol | Fold change | P value | Function | Protein expression MDA MBA231 relative to JL-Contra-3 | Protein expression MCF7 relative to JL-Contra-3 |
| --- | --- | --- | --- | --- | --- | --- |
| 1. | SFRS3 | 1.334 (↑) | 0.001 | SFRS3 (serine and arginine-rich splicing factor 3); involved in the regulation of alternative splicing of pre-mRNA molecules | ↑ | ↑* |
| 2. | BAG3 | 1.401 (↑) | 0.015 | Bcl-2-associated gene 3; BAG family of co-chaperones that interact with HSP70 (heat shock protein 70) | ↓ | ↓ |
| 3. | DDX5 | 1.479 (↑) | 0.026 | DEAD-box helicase 5; pre-mRNA splicing, mRNA export, translation, and decay | ↑ | ↑* |
| 4. | SNRPD3 | 1.616 (↑) | 0.027 | Small nuclear ribonucleoprotein D3; component of the spliceosome | ↑ | ↑* |
| 5. | PLEC1 | 1.946 (↑) | 0.000001 | Plectin 1; cell structure and organization, as well as in signaling pathways and cellular functions such as cell motility, adhesion, and survival | ↑* | ↓ |
| 6. | ATP5A1 | 2.028 (↑) | 0.039 | ATP synthase subunit alpha; responsible to produce ATP (adenosine triphosphate) in the mitochondria of cells, been suggested to play a role in cancer cell metabolism and survival | ↑ | ↑* |
| 7. | HSPB1 | 2.060 (↑) | 0.023 | Heat Shock Protein Family B Member 1, aka HSP27; induced in response to various cellular stresses, including heat shock, oxidative stress, and exposure to toxins or radiation | ↓ | ↑* |
| 8. | PRSS1 | 2.152 (↑) | 0.048 | Serine Protease I (trypsin); promotes cancer cell proliferation, invasion, etc. | ↓ | ↑ |
| 9. | DSP | 2.159 (↑) | 0.003 | Desmoplakin; obligate component of functional desmosomes, inactivated in cancer | ↑ | ↑ |
| 10. | ANXA1 | 2.290 (↑) | 0.014 | Annexin A1 is a member of the annexin family of calcium-dependent phospholipid-binding proteins, role in inflammation, cell differentiation, and apoptosis | ↑ | ↓* |
| 11. | PDAP1 | 2.402 (↑) | 0.037 | Platelet-derived growth factor (PDGF); multifunctional protein that plays a role in several cellular processes, including DNA replication, repair, and apoptosis, increased in cancer, might be specific to DCIS | ↓ | ↓* |
| 12. | KRT1 | 2.960 (↑) | 0.011 | Keratin 1; structural protein major component of the cytoskeleton in epithelial cells. | ↑ | ↓ |
| 13. | SFRS7 | 2.964 (↑) | 0.012 | SFRS7 (Splicing factor, arginine/serine-rich 7); involved in pre-mRNA splicing, regulation of alternative splicing, involved in the splicing of several genes related to cancer: CD44, BRCA1, and BCL-X. Dysregulation of SFRS7 expression and splicing activity has been linked to various cancers | ↑ | ↑* |
| 14. | RPL11 | 3.669 (↑) | 0.021 | ribosomal protein that is a component of the 60S subunit of the ribosome, implicated in cancer, interacts with MDM2 | ↑ (0.07) | ↓ |
| 15. | RPL7A | 3.951 (↑) | 0.049 | ribosomal protein; plays a regulatory role in the translation apparatus, implicated in cancer | ↑ | ↓ |
| 16. | RPL31 | 4.000 (↑) | 0.018 | ribosomal protein; In addition to its role in protein synthesis, RPL31 has also been implicated in cancer development and progression. Studies have shown that RPL31 expression is elevated in prostate, gastric and colorectal cancers | ↑ | ↑ |
| 17. | TPM1 | 4.333 (↑) | 0.012 | Tropomyosin 1; mechanical integrity of the cell, regulating cell shape, adhesion, and migration, may promote tumor invasion by downregulating Ecadherin and upregulating MMPs | ↑ | ↓ |
| 18. | MECP2 | 4.663 (↑) | 0.002 | Methyl-CpG-binding protein 2; binds to methylated DNA and plays a role in silencing genes that are not needed by a particular cell type | ↑* | ↓ |
| 19. | RPL8 | 5.462 (↑) | 0.002 | Ribosomal Protein L8; protein component of the ribosome, role in the binding and positioning of tRNAs during translation | ↑ | ↓ |
| 20. | ALB | 6.944 (↑) | 0.003 | Albumin; transports hormones, fatty acids, etc. | ↑ | ↓ |
| 21. | CKAP4 | 7.550 (↑) | 0.0002 | Cytoskeleton-Associated Protein 4; crucial role in the organization and maintenance of the cellular ER, implicated in cell proliferation and migration, vesicle transport | ↑* | ↓* |

Using Metascape (metascape.org), we determined pathways represented by downregulated proteins (JL-DCIS-3 relative to JL-Contra-3) including microRNAs associated with cancer, regulation of RNA splicing, apoptotic execution phase, and regulation of intrinsic apoptotic signaling, all of which have been associated with cancer (**Fig. 1A**). In contrast, the upregulated proteins in JL-DCIS-3 vs. JL-Contra-3 represented pathways that included locomotion (regulation of blood vessel endothelial cell migration), metabolism of RNA, peptide cross linking, and other pathways (**Fig. 1B**). Spliceosomes were significant in both upregulated and downregulated protein groups.

**Fig. 1A.**
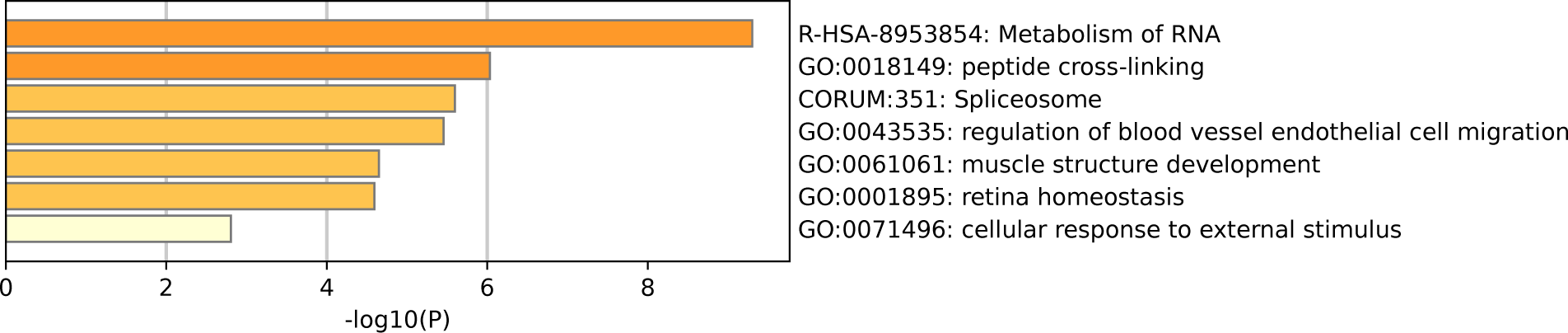
Enrichment heat map generated by Metascape showing pathways represented by upregulated proteins in JL-DCIS-3 relative to JL-Contra-3.

**Fig. 1B.**
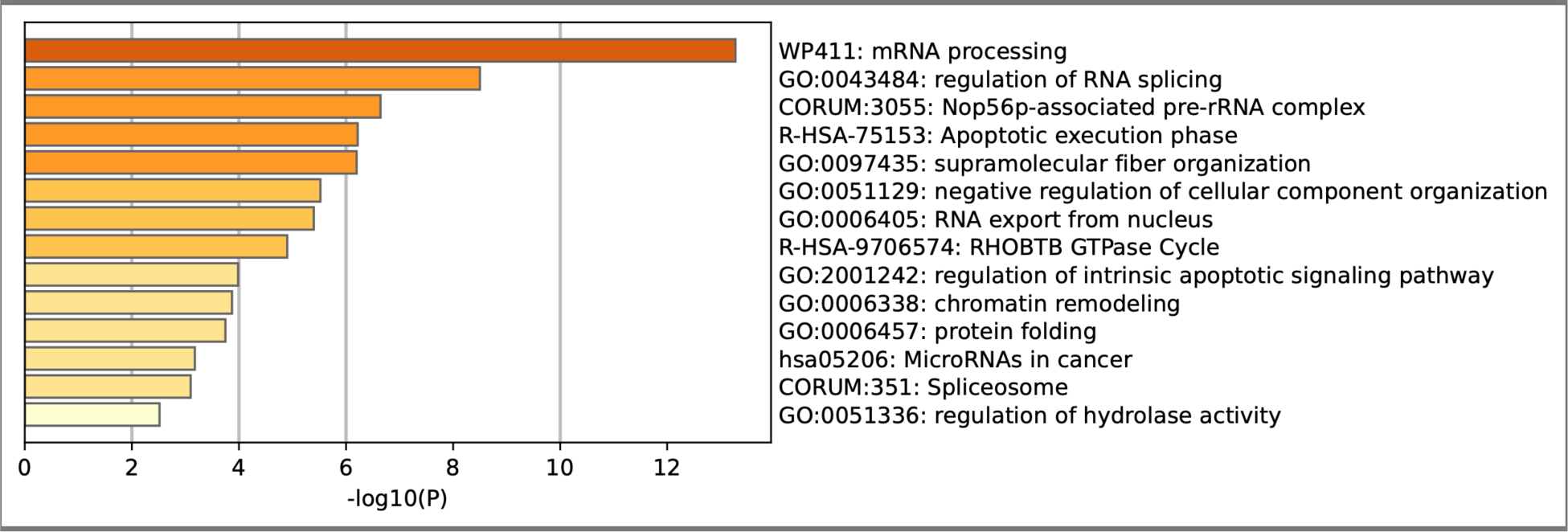
Enrichment heat map generated by Metascape showing pathways represented by downregulated proteins in JL-DCIS-3 relative to JL-Contra-3.

### Multi-omic analysis of JL-DCIS-3 vs. JL-Contra-3

We then specifically analyzed the mRNA expression of the 60 differentially expressed proteins in JL-DCIS-3, JL-Contra-3, and a non-diseased breast reduction mammoplasty cell line (JL-BRL-6) and included RNAs extracted from additional patient-derived breast cancer cultures that were established by the same culture system (Latimer, 2000b; Latimer et al., 2010) from stages I, II, III, and IV breast tumors. Also included were RNAs from the commercially available MCF-7 and MDA-MB231 breast cancer cell lines because they are literature standards. This mRNA analysis strategy was chosen so that a molecular test driven by protein expression could be developed in the future.

The genes of interest were selected using the following criteria. First, RNA analyses should reveal a significant difference between DCIS and contralateral breast, consistent with the proteomic analysis. Second, there should be no significant difference in the expression of the gene of interest between JL-Contra-3 and the independent, unrelated breast reduction mammoplasty PDCEE JL-BRL-6, a cell line that serves as a well-studied standard in our laboratory (Visus et al., 2011).

We identified 10 NMPs that were significantly differentially expressed in JL-DCIS-3 relative to JL-Contra-3 in terms of both protein and RNA expression levels (**Fig 2 and 3**). In addition, these NMP genes had similar mRNA expression in JL-Contra-3 and JL-BRL-6. Since JL-Contra-3 and JL-BRL-6 both have a normal female karyotypes and were generated using the same methodology, this was an independent validation of the non-diseased nature of the contralateral breast from the DCIS patient from which these cell lines were derived. This does not imply that all women with DCIS have non-diseased contralateral breast tissue.

**Fig. 2.**
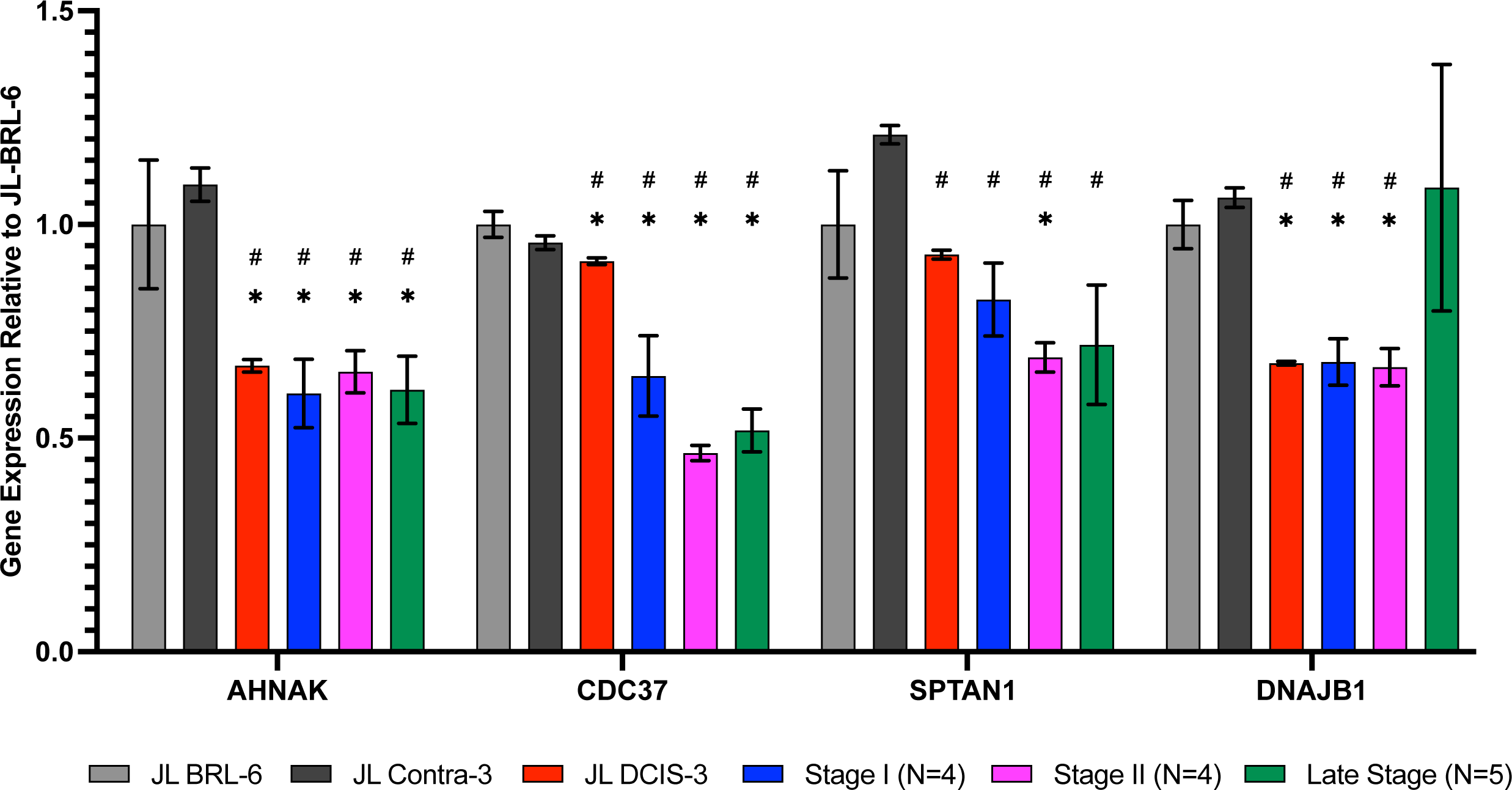
RNA expression of nuclear matrix protein genes identified as significantly downregulated in proteomics analysis for JL DCIS-3 (red bar) and BC stages (blue, pink, and green bars) relative to non-diseased breast reduction (JL-BRL-6) patient derived cells (gray bar) and to JL-Contra-3 (black bar). These genes manifest no significant difference between JL-BRL-6 (gray bar) and JL-contralateral-3 (black bar; p> 0.05). Asterisk (*) denotes that significant different relative to JL-BRL-6. Pound (#) denotes that significant different relative to JL-Contra-3.

**Fig. 3.**
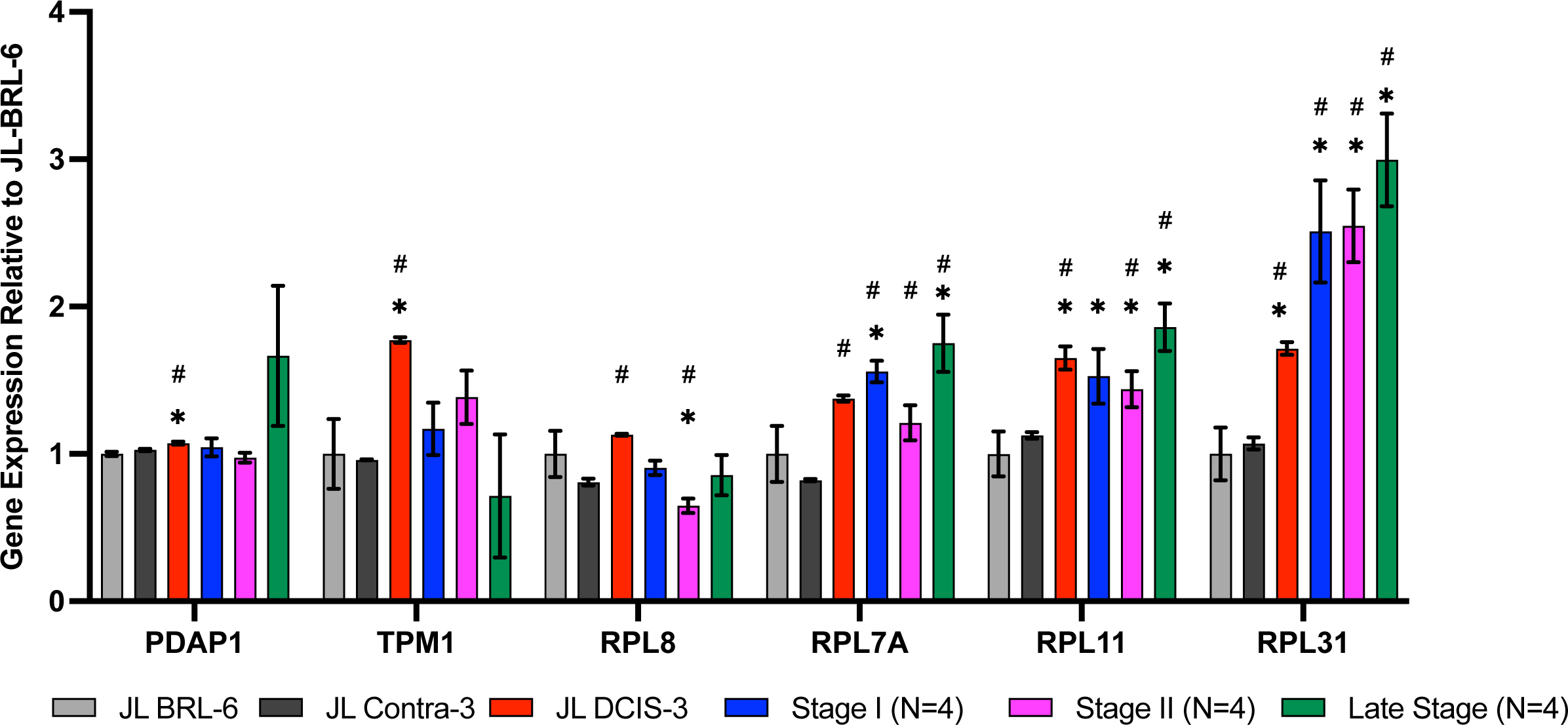
RNA expression of nuclear matrix protein genes identified as significantly upregulated in proteomics analysis for JL DCIS-3 (red bar) and BC stages (blue, pink, and green bars) relative to patient derived non-diseased breast reduction cells (gray bar) and JL-Contra-3 (black bar). These genes manifest no significant difference between JL-Contra-3 (black bar) and JL-BRL-6 (gray bar; p > 0.05). Asterisk (*) denotes that significant different relative to JL-BRL-6. Pound (#) denotes that significant different relative to JL-Contra-3.

Of the 10 differentially expressed genes in the DCIS and contralateral isogenic pair, four of ten genes were downregulated in the DCIS sample relative to the non-diseased contralateral sample (**Fig 2**). These four genes, AHNAK (p < 0.01) (neuroblast differentiation-associated protein AHNAK; aka desmoyokin), CDC37 (p = 0.03) (leukocyte antigen CD37), SPTAN1 (p < 0.01) (spectrin alpha, non-erythrocytic 1), and DNAJB1 (p < 0.01) (DnaJ heat shock protein family [Hsp40] member B1), were also significantly downregulated in invasive BC. This suggests that JL-DCIS-3 manifests some of the abnormal characteristics of invasive BC stages. All four of these genes are implicated in cancer (Lee et al., 2014; Cui et al., 2015; Ackermann et al., 2019; Ono et al., 2020).

JL-DCIS-3 gene expression was significantly upregulated in 6 of 10 NMP genes relative to non-diseased JL Contra-3: RPL31(ribosomal protein L31; p = 0.001), RPL-11 (ribosomal protein L11; p < 0.001), RPL7A (ribosomal protein 7A; p < 0.01), PDAP1 (PDGFA associated protein 1; p = 0.01), TPM1 (tropomyosin 1; p < 0.001), and RPL8 (ribosomal protein L8; p < 0.001; **Fig 2**). Ribosome biogenesis occurs in the nucleolus and involves the action of 80 ribosomal proteins (RPs), four ribosomal RNAs (rRNAs) other associated proteins, and small nucleolar RNAs (snoRNAs) (El Khoury & Nasr, 2021). Several ribosomal proteins are overexpressed in human cancers including prostate, lung, breast, pancreatic, liver and colon (Goudarzi & Lindström, 2016; Penzo et al., 2019). Gene expression of PDAP1 (PDGF, Platelet Derived Growth Factor), a multifunctional protein, is increased in cancer and plays a role in DNA replication, repair, and invasion (Choi et al., 2011; H. Cui et al., 2022).

### NMP Markers Associated with BC Progression

The 10 genes that were significantly differentially expressed in JL-DCIS-3 relative to JL-Contra-3 in both protein and steady state RNA were also associated with changes in the progression of BC (**Fig 2 and 3**). These are good candidates for inclusion in a molecular test panel for aggressive DCIS.

The protein levels of AHNAK, CDC37, and SPTAN1 were significantly downregulated in JL-DCIS-3 vs JL-Contra-3 but were also significantly downregulated in the late-stage BC cell lines MDA-MB231 and MCF-7 relative to JL-Contra-3 (AHNAK [p <0.00001 for both], CDC37 [p = 0.009, 0.006 respectively], SPTAN1 [p <0.00001 for both]). DNAJB1 gene expression was significantly downregulated in early-stage BC compared with non-diseased breast cells (p < 0.01) and was not significantly upregulated in the late-stage BC compared to JL-Contra-3 and JL-BRL-6 (p = 0.4). Conversely, the protein level was downregulated in MDA-MB231 and MCF-7 cells (p = 0.02 and 0.14 respectively). AHNAK and CDC-37 were significantly downregulated in JL-DCIS-3 as well as in all invasive BC stages relative to non-diseased contralateral JL-DCIS-3 and JL-BRL-6 (all p < 0.04) (**Fig 2**, **Table 1**). Linear regression analysis showed a strong association between disease progression and RNA expression of CDC37, SPTAN1, RPL7A, RPL11 and RPL31 (p < 0.05). These genes were upregulated as both protein and mRNA in JL-DCIS-3 relative to JL-Contra-3.

Spectrins are a family of filamentous cytoskeletal structures that function as scaffold proteins. They stabilize the plasma membrane and arrange organelles. SPTAN1 is implicated in DNA repair and cell cycle regulation (Lambert, 2018, 2019). Higher protein and mRNA levels of SPTAN1 are associated with longer patient survival times in colon cancers. However, MLH1-deficient colorectal cancers exhibit reduced levels of this cytoskeleton scaffold gene (SPTAN1) (Ackermann et al., 2019). Our DCIS data agree with the latter study.

The expression of the RPL31 gene increased in JL-DCIS-3 relative to JL-Contral-3 but also increased with stage progression (**Fig 2**). However, the protein level of RPL31 in MDA-MB231 and MCF-7 cells was also upregulated relative to JL-Contra-1, but not significantly so (p = 0.14 and 0.24, respectively).

RPL11 was significantly upregulated in JL-DCIS-3, stage II and late-stage invasive BC relative to non-diseased JL-BRL-6 and JL-Contra-3 (p <0.05). RPL11 expression was also upregulated in stage I BC relative to JL-BRL-6 (p = 0.04) and JL-Contra-1 (p= 0.06). Similarly, RPL7A gene expression was significantly upregulated in JL-DCIS-1 and invasive breast cancer stages relative to non-diseased JL-Contra-1 (p <0.02), and only stage I and stage IV invasive breast cancer were significantly upregulated relative to non-diseased JL-BRL-6 (p = 0.01 and 0.02 respectively; **Fig 2**).

RPL8 was significantly upregulated in JL-DCIS-3 relative to its isogenic matched contralateral in both protein and gene expression levels (p < 0.01). A trend in upregulation was also observed in JL-DCIS-3 relative to non-diseased JL-BRL-6, but was not significant (p = 0.22; **Fig 2**). Furthermore, there was variation in RPL8 gene expression between invasive BC stages relative to non-diseased cultures (**Fig 2**).

PDAP1 and TPM1 were significantly upregulated in JL-DCIS-3 relative to non-diseased breast reduction and JL Contra-1 (p < 0.02). However, PDAP1 and TPM1gene expression was not significantly different between invasive BC stages and non-diseased cells (p > 0.05; **Fig 2**). This may indicate that JL-DCIS-3 as an individual tumor-derived culture simply does not adhere to patterns of expression for these genes as seen in invasive cancers. These genes could also be DCIS-specific biomarkers.

### Genes with Opposite Patterns in Protein and Steady State RNA Expression

We also identified 13 genes that were significantly differentially expressed between JL-Contra-3 and JL-DCIS-3 in both RNA sequencing and proteomic results, but the RNA expression changes were not consistent with the protein levels found by mass spectrometry (**Fig 4**). HSPB1 (heat shock protein beta-1) was significantly downregulated in RNA expression (p value = 0.044) but the protein level was significantly elevated (p = 0.023).

**Fig. 4.**
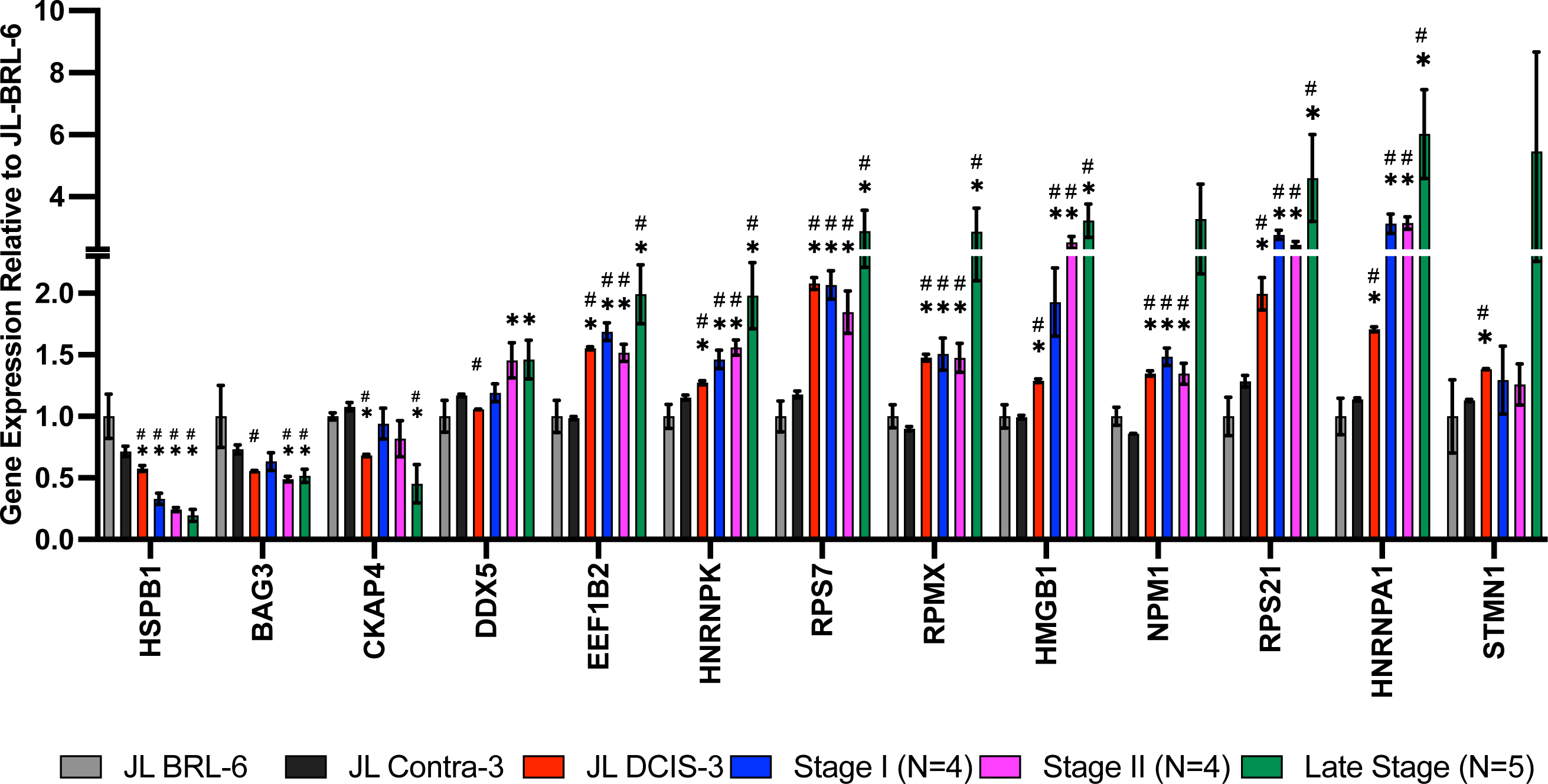
mRNA expression from RNA sequencing data of NMPs that were among the 60 proteins whose levels were significantly different between JL-DCIS-3 and JL-Contra-3 patient-derived cells. The RNA expression for these genes showed the opposite trend compared with the proteomics analysis.

Interestingly, the RNA expression of HSPB1 decreased with stage progression (**Fig 4**). Small heat shock proteins like HSPB1 are associated with multiple processes in cancer including invasion, control of apoptosis and drug resistance (Xiong et al., 2020). In contrast, the RNA expression of EEF1B2 (eukaryotic translation elongation factor 1 beta 2; p < 0.0001), HMGB1 (high mobility group box 1; p < 0.0001), and HNRNPA1 (heterogeneous nuclear ribonucleoprotein A1; p = 0.0003) were all significantly upregulated in JL-DCIS-3 relative to JL-Contra-3, and the RNA expression was increased with tumor stage progression. Even though the RNA levels of HSPB1, EEF1B2, HMGB1, HNRNPA1, RPS7 (ribosomal protein S7), RPS21 (ribosomal protein S21), and HNRNPK (heterogenous nuclear ribonucleoprotein K) did not correlate with protein levels, JL-DCIS-3 was consistent with the invasive breast cancer pattern (all stages) in the mRNA expression of these genes (**Fig 4**). Overexpression of EEF1B2 and HNRNPA1 has been previously linked with cancer progression and poor prognosis in breast and lung cancer (Hassan et al., 2018; Jia et al., 2019; Roy et al., 2017). Similarly, the overexpression of cytoplasmic HMGB1 stimulates tumor cell proliferation and metastasis (Yuan et al., 2020). Conversely, nuclear HMGB1 is implicated in maintaining genomic integrity, and losing nucleus HMGB1 results in genomic instability and carcinogenesis (Wang and Zhang, 2020). This discrepancy in HMGB1 function according to cellular location could potentially account for differences between protein and RNA expression, as our study focused on the proteomics of the cellular nuclear matrix portion.

## Discussion

DCIS offers a critical insight into early BC that may or may not progress to clinical disease. The determination of indolent versus aggressive DCIS is of paramount importance because all DCIS cases are treated, and the treatments can cause permanent complications, for example, damage to the underlying lung from radiation.

Proteomic-transcriptomic strategies are powerful for determining biomarkers of cancer that may be useful clinically at different levels. Historically, biomarker identification at the level of proteins using immunohistochemistry (IHC) has been an important standard for cancer diagnosis. More recently, RNA biomarkers have become useful in the form of prognostic tests such as Oncotype DX and MammaPrint for breast cancer prognosis. With the advent of tumor sequencing, mutational burden in general, as well as specific druggable mutations, have been added to the assessment of cancer prognosis and treatment tailoring.

The morphological appearance of the nucleus, the nucleoli, and the nucleus-to-cytoplasmic ratio represent key features used to identify tumor cells and are considered in the determination of nuclear grade (Zink et al., 2004). In this study, we focused on the nucleus by specifically analyzing alterations in the expression of NMPs. In this way, we have not diluted our mass spectrometric proteomic findings with the additional abundant proteins located in the cytoplasm. We used our proteomic findings to prioritize the most nucleus-centric RNA findings (using deep RNA sequencing).

We used as model systems a novel stage 0 patient-derived BC culture (JL-DCIS-3) and compared it to an isogenic non-diseased contralateral culture (JL-Contra-3). This is the most direct comparison that can be made, and, although it represents one isogenic set of comparisons within a single individual, it was contextually expanded to include comparisons to multiple patient-derived cultures and cell lines from tumors of stages I, II, III and IV. Our hypothesis was that changes within one patient’s tissues will reveal early alterations linked to the onset of disease for that patient that may be generalizable to other cases of DCIS. Surprisingly, many of the RNA changes we saw in DCIS were also consistent with disease progression. Looked at another way, it was surprising that so much of what we know about disrupted pathways in advanced cancer were identified very early in disease progression (mRNA processing, spliceosomes, metabolic differences). This was not always the case, with some NMP-related genes showing opposite trends at the protein vs. RNA levels (**Fig 4**). In addition, since RNA sequencing was performed on whole cell cultures (JL-DCIS-3 vs JL-Contra-DCIS-3) and because it was not possible to isolate RNA after NMP protocols, sometimes the matching RNA expression could not be detected for comparison with mass spectrometric analyses.

This study is unique in that it was driven by NMP differences using the growing compartment of a DCIS and matching isogenic contralateral culture. This constitutes a more functional preparation of related proteins. Most studies on DCIS biomarkers are focused on tissue or laser-captured whole DCIS cells. The impact of biomarkers in the nuclear matrix structure within the nucleus would therefore be diluted in these laser capture comparisons because in the latter preparation the entire cell is being utilized, including all of the cytoplasmic proteins in addition to those from the nucleus.

The major lesson from this study may be that focusing on a specific functional compartment of the cell allows for the identification of biomarkers of DCIS that will not emerge from harvesting the entire cell and looking for protein or mRNA differences. What is also unusual in this work is the integration of proteomics with RNA sequencing analyses to determine biomarkers that follow the same patterns at both levels. This does not diminish the importance of proteins that show trends that are opposite of the mRNA due to differences in post transcriptional processing, but these are the subject of another biomarker group. In fact, if differences exist between protein and RNA in this study, it could be an indication that this protein exists in significant quantities outside the nuclear matrix. What we measured with mass spectrometry was specific to the nuclear matrix.

Using mass spectrometry on nuclear matrix preparations, we identified 60 proteins whose levels were significantly different between JL-DCIS-3 and JL-Contra-3. After identifying nuclear matrix protein whose expression were significantly different between JL-DCIS-3 and JL-Contra-3, we profiled the gene expression of those 60 differing NMPs in JL-DCIS-3, JL-Contra-3, a non-diseased breast reduction culture (JL-BRL-6) and four stage I, four stage II, and four late-stage breast cancer patient-derived cultures or commercially available cell lines. Ten of 60 NMP proteins were similarly expressed between non-diseased breast (JL-BRL-6 and JL-Contra-3) but were significantly different in JL-DCIS-3. We identified 10 NMPs that were significantly differentially expressed in JL-DCIS-3 relative to JL-Contra-3 cultures in terms of both protein and gene expression levels (**Fig 1 and 2**). Of the 10 genes, six genes *were significantly different* in both JL-DCIS-3 and invasive breast cancer cultures compared with non-diseased cultures. These six genes were AHNAK, CDC37, DNAJB1, RPL7A, RPL11, and RPL31.

AHNAK protein was downregulated in JL-DCIS-3 relative to JL-Contra-3 by 5.4-fold. Downregulation of AHNAK has previously been observed in ovarian cancer (Cai et al., 2021), lung cancer (Park et al., 2018), brain tumor (Zhao et al., 2017) and BC (Lee et al., 2014). In this study, we saw downregulation of AHNAK mRNA expression for all BC stages as well as in both mRNA and protein expressions of JL-DCIS-3, MDA-MB231, and MCF-7. AHNAK mRNA overexpression has been found to inhibit triple negative BC cell growth and lung metastasis in **in vivo** xenografts (Chen et al., 2017). Since we observed a reduction of AHNAK mRNA and protein in both early and late-stage BC, AHNAK could be a reliable prognostic indicator for DCIS patients to distinguish between indolent and aggressive DCIS, presumably representing aggressive disease.

The protein level of CDC37 was reduced more than 9-fold in JL-DCIS-3 relative to JL-Contra-3. CDC37 plays an important role in proliferation and transformation of tumor cells by maintaining protein kinase activity. CDC37 has a critical role in progression of oral (Ono et al., 2020) and prostate cancer (Eguchi et al., 2019). Several studies have shown the importance of targeting CDC37 for cancer therapy (Smith et al., 2009). However, in this study, we found the CDC37 protein and RNA levels were lower in diseased JL-DCIS-3 relative to non-diseased JL-Contra-3 and JL-BRL-6. The reduction was also maintained in gene expression of invasive BC, stage I through stage IV (**Fig 1**). This may indicate that the role of CDC37 in BC is different, perhaps acting as a tumor suppressor gene rather than an oncogene.

DNAJB1 encodes a member of the DNAJ or HSP40 family of proteins. These genes function as one of the two major classes of molecular chaperones involved in protein folding and oligomeric protein assembly, as well as in the proteosome pathway, endoplasmic reticulum stress and viral infection. Previous studies have shown that DNAJB1 has a critical role in both tumor suppression and progression based on the activity of p53 in different cancers (Cui et al., 2015; Qi et al., 2014). Cui et al. (2015) showed that DNAJB interacted with PDC5 in a lung cancer cell line and inhibited the apoptotic function of p53. Conversely, Qi et al. (2014) showed that DNAJB1 activates p53 through stabilizing MDM2, a major ubiquitin ligase that inhibits p53. Inhibition of DNAJB1 increased the proliferation and tumor growth of MCF-7 cells (Qi et al., 2014). The latter work was consistent with our findings.

The protein level of DNAJB1 was four-fold lower in diseased JL-DCIS-3 relative to non-diseased JL Contra-1 in the nuclear matrix. The reduction was also shown by mRNA gene expression analysis in JL DCIS-3 and early-stage BC. In late-stage BC, the protein level of DNAJB1 was reduced by 3.9-fold in MCF-7 (although not significantly) and significantly reduced by 2.5-fold in MDA-MB231. However, we did not see a reduction in DNAJB1 RNA expression in late-stage BC (**Table 1**, **Fig 2**).

Ribosome biogenesis occurs in the nucleolus. Multiple ribosome biogenesis proteins were significantly differentially expressed in our isogenic pair of PDCs. RPL11, RPL31 and RPL7A protein levels were higher and mRNA expression levels were upregulated in diseased JL-DCIS-1 relative to non-diseased JL-Contra-1 (**Table 2**). Similarly, the mRNA expression of RPL11, RPL31, and RPL7A were upregulated in invasive BC (**Fig 2**). RPs are essential in ribosome biogenesis, and they are synthesized in the cytoplasm then imported into the nucleus to assemble with rRNA. RPs also have extra-ribosomal functions in cellular processes, including cell migration and invasion (Liu et al., 2007; Yang et al., 2013), differentiation (Da Costa et al., 2003; Zhan et al., 2010), and DNA repair (Kim et al., 1995; Wang et al., 2015). Changes in RP expression have previously been used as prognostic or predictive indicators to distinguish between normal and cancer cells (Goudarzi and Lindström, 2016). For example, high RPL19 expression is associated with poor survival in prostate cancer (Bee et al., 2006). In contrast, a defect in ribosome biogenesis increases cancer susceptibility (Narla & Ebert, 2010; Ruggero & Pandolfi, 2003). RPs can act as tumor suppressor or tumor promoting genes based on their effect on mRNA translation of their binding partner, either oncogenes or tumor suppressor genes (Goudarzi and Lindström, 2016). RPL7A has been found to be overexpressed in prostate and colorectal cancer (Vaarala et al., 1998). In contrast, downregulation of RPL7A is associated with poor prognosis of overall survival of osteosarcoma patients with lung metastasis (Yao et al., 2009). In our findings, RPL7A was significantly overexpressed in JL-DCIS-1 relative to JL-Contra-1 in both protein and gene expression levels as well as in all invasive BC stages.

RPL31 is involved in cell proliferation and has been found to be overexpressed in colon cancer (de las Heras-Rubio et al., 2014; W. Wang et al., 2015). In our findings, RPL31 was significantly higher in both protein and gene expression levels of JL-DCIS-1 relative to non-diseased cells (**Table 2**). The gene expression of RPL31 was significantly higher in all invasive BC samples (**Fig 2**). The protein level was four-fold higher in diseased JL-DCIS-1 relative to non-diseased JL-Contra-1 (**Table 1**) and 1.5-fold higher in stage IV, while the gene expression level of RPL31 was expressed highly in early and late-stage BC compared with non-diseased cells. That may indicate that high translation efficiency of RPL31 mRNA in pre-invasive BC (DCIS) was decreased in late-stage BC. RPL31 may be a stage-specific protein. Specifically, the protein level of RPL31 could be an indicator of diseased DCIS (stage 0).

Upregulation of RPL11 has associated with activation of the p53 pathway and inhibition of cell proliferation through stabilization of E2F1(Bhat et al., 2004; Dai et al., 2008). Moreover, the anti-proliferative effect of RPL11 negatively regulates c-Myc levels and activity (Dai et al., 2010). We found both protein and gene expression levels of RPL11 were significantly upregulated in JL-DCIS-1 relative to JL-Contra-1. Similarly, the gene expression of late-stage BC was significantly upregulated compared to JL-BRL-6 (**Fig 2**). However, these results were not significant in stages I and II.

The protein level of RPL11 was insignificantly lower in MCF-7 and only 1.5-fold higher in MDA-MB231 cells. The change in protein level of RPL11 between DCIS and late-stage BC may indicate the involvement of RPL11 in BC progression. It might not be a good indicator for early stage BC but could be targeted in late stages.

CKAP4 and HSPB1 protein levels were significantly upregulated, by seven- and two-fold, respectively, in diseased JL-DCIS-3 tissue compared to non-diseased contralateral tissue (**Table 2**). However, the gene expression of CKAP4 was significantly downregulated in JL-DCIS-3 and late-stage BC, while HSPB1 gene expression decreased with stage progression (**Fig 4**). A recent study found that CKAP4 enhances cell invasiveness and migration (Osugi et al., 2019). CKAP4 gene expression was not significantly different between JL-DCIS-3 and late-stage BC, while a five-fold difference in protein level between JL-DCIS-3 and late-stage BC was observed. This difference in protein levels between pre-invasive DCIS and late-stage BC might suggest that CKAP4 may play an important role in the etiology of pre-invasive DCIS. Similarly, HSPB1 protein was significantly two-fold higher in JL-DCIS-3 as well as in MCF-7 relative to non-diseased contralateral tissue. However, no significant difference in HSBP1 protein level was observed in MDA-MB231(**Table 2**). Conversely, the gene expression of HSPB1 was downregulated with stage progression (**Fig 4**). HSPB1 has been suggested to be involved in cell proliferation (Treweek et al., 2015) and inhibition of apoptosis in BC (Carra et al., 2019).

In conclusion, we identified 60 NMPs whose protein levels are significantly different between matching isogenic diseased DCIS and non-diseased contralateral normal breast epithelial tissue. The alterations in these NMPs could be involved in DCIS formation and could be helpful in diagnosis of DCIS cases. By profiling the gene expression of these NMPs, we identified 10 genes in diseased JL-DCIS-3 that were significantly different compared to non-diseased contralateral and breast reduction tissues. This suggests that these 10 NMPs might be linked to progression of DCIS into invasive BC and could be helpful in defining invasive versus indolent DCIS.

## Acknowledgments

This work was supported in part by grants from the U.S. National Institutes of Health (1T32EB001026), the U.S. Department of Defense (W81XWH-0701-0660) and Susan G. Komen For The Cure (DISP0707276).

## References

Ackermann, A., Schrecker, C., Bon, D., Friedrichs, N., Bankov, K., Wild, P., Plotz, G., Zeuzem, S., Herrmann, E., Hansmann, M.-L., & Brieger, A. (2019). Downregulation of SPTAN1 is related to MLH1 deficiency and metastasis in colorectal cancer. PLOS ONE, 14(3), e0213411. 10.1371/journal.pone.0213411

Bee, A., Ke, Y., Forootan, S., Lin, K., Beesley, C., Forrest, S. E., & Foster, C. S. (2006). Ribosomal protein L19 is a prognostic marker for human prostate cancer. Clinical Cancer Research, 12(7), 2061 LP – 2065. 10.1158/1078-0432.CCR-05-2445

Bhat, K. P., Itahana, K., Jin, A., & Zhang, Y. (2004). Essential role of ribosomal protein L11 in mediating growth inhibition-induced p53 activation. The EMBO Journal, 23(12), 2402– 2412. 10.1038/sj.emboj.7600247

Cai, Y., Hu, Y., Yu, F., Tong, W., Wang, S., Sheng, S., & Zhu, J. (2021). AHNAK suppresses ovarian cancer progression through the Wnt/β-catenin signaling pathway. Aging, 13(20), 23579–23587. 10.18632/aging.203473

Carra, S., Alberti, S., Benesch, J. L. P., Boelens, W., Buchner, J., Carver, J. A., Cecconi, C., Ecroyd, H., Gusev, N., Hightower, L. E., Klevit, R. E., Lee, H. O., Liberek, K., Lockwood, B., Poletti, A., Timmerman, V., Toth, M. E., Vierling, E., Wu, T., & Tanguay, R. M. (2019). Small heat shock proteins: multifaceted proteins with important implications for life. Cell Stress & Chaperones, 24(2), 295–308. 10.1007/s12192-019-00979-z

Chen, B., Wang, J., Dai, D., Zhou, Q., Guo, X., Tian, Z., Huang, X., Yang, L., Tang, H., & Xie, X. (2017). AHNAK suppresses tumour proliferation and invasion by targeting multiple pathways in triple-negative breast cancer. Journal of Experimental & Clinical Cancer Research, 36(1), 65. 10.1186/s13046-017-0522-4

Choi, S., Jang, J. H., & Kim, K. R. (2011). Analysis of differentially expressed genes in human rectal carcinoma using suppression subtractive hybridization. Clinical and Experimental Medicine, 11(4), 219–226. 10.1007/s10238-010-0130-5

Collins, L. C., Tamimi, R. M., Baer, H. J., Connolly, J. L., Colditz, G. a., & Schnitt, S. J. (2005). Outcome of patients with ductal carcinoma in situ untreated after diagnostic biopsy: Results from the nurses’ health study. Cancer, 103(March), 1778–1784. 10.1002/cncr.20979

Cui, H., Wei, W., Qian, M., Tian, R., Fu, X., Li, H., Nan, G., Yang, T., Lin, P., Chen, X., Zhu, Y., Wang, B., Sun, X., Dou, J., Jiang, J., Li, L., Wang, S., & Chen, Z. (2022). PDGFA-associated protein 1 is a novel target of c-Myc and contributes to colorectal cancer initiation and progression. Cancer Communications, 42(8), 750–767. 10.1002/cac2.12322

Cui, X., Choi, H. K., Choi, Y. S., Park, S. Y., Sung, G. J., Lee, Y. H., Lee, J., Jun, W. J., Kim, K., Choi, K. C., & Yoon, H. G. (2015). DNAJB1 destabilizes PDCD5 to suppress p53-mediated apoptosis. Cancer Letters, 357(1), 307–315. 10.1016/j.canlet.2014.11.041

Da Costa, L., Narla, G., Willig, T.-N., Peters, L. L., Parra, M., Fixler, J., Tchernia, G., & Mohandas, N. (2003). Ribosomal protein S19 expression during erythroid differentiation. Blood, 101(1), 318–324. 10.1182/blood-2002-04-1131

Dai, M.-S., Sun, X.-X., & Lu, H. (2008). Aberrant expression of nucleostemin activates p53 and induces cell cycle arrest via inhibition of MDM2. Molecular and Cellular Biology, 28(13), 4365–4376. 10.1128/MCB.01662-07

Dai, M.-S., Sun, X.-X., & Lu, H. (2010). Ribosomal protein L11 associates with c-Myc at 5 S rRNA and tRNA genes and regulates their expression. The Journal of Biological Chemistry, 285(17), 12587–12594. 10.1074/jbc.M109.056259

de las Heras-Rubio, A., Perucho, L., Paciucci, R., Vilardell, J., & LLeonart, M. E. (2014). Ribosomal proteins as novel players in tumorigenesis. Cancer and Metastasis Reviews, 33(1), 115–141. 10.1007/s10555-013-9460-6

Dellaire, G. (2003). The Nuclear Protein Database (NPD): sub-nuclear localisation and functional annotation of the nuclear proteome. Nucleic Acids Research, 31(1), 328–330. 10.1093/nar/gkg018

Doke, K., Butler, S., & Mitchell, M. P. (2018). Current therapeutic approaches to DCIS. Journal of Mammary Gland Biology and Neoplasia, September, 279–291.

Duffy, S. W., Tabár, L., Yen, A. M.-F., Dean, P. B., Smith, R. A., Jonsson, H., Törnberg, S., Chen, S. L.-S., Chiu, S. Y.-H., Fann, J. C.-Y., Ku, M. M.-S., Wu, W. Y.-Y., Hsu, C.-Y., Chen, Y.-C., Svane, G., Azavedo, E., Grundström, H., Sundén, P., Leifland, K., … Chen, T. H.-H. (2020). Mammography screening reduces rates of advanced and fatal breast cancers: Results in 549,091 women. Cancer, 126(13), 2971–2979. 10.1002/cncr.32859

Eguchi, T., Prince, T. L., Tran, M. T., Sogawa, C., Lang, B. J., & Calderwood, S. K. (2019). MZF1 and SCAND1 reciprocally regulate CDC37 gene expression in prostate cancer. Cancers, 11(6), 792. 10.3390/cancers11060792

El Khoury, W., & Nasr, Z. (2021). Deregulation of ribosomal proteins in human cancers. Bioscience Reports, 41(12). 10.1042/BSR20211577

Erbas, B., Provenzano, E., Armes, J., & Gertig, D. (2006). The natural history of ductal carcinoma in situ of the breast: A review. Breast Cancer Research and Treatment, 97(2), 135–144. 10.1007/s10549-005-9101-z

Goudarzi, K. M., & Lindström, M. S. (2016). Role of ribosomal protein mutations in tumor development (Review). International Journal of Oncology, 48(4), 1313–1324. 10.3892/ijo.2016.3387

Hassan, Md. K., Kumar, D., Naik, M., & Dixit, M. (2018). The expression profile and prognostic significance of eukaryotic translation elongation factors in different cancers. PLOS ONE, 13(1), e0191377. 10.1371/journal.pone.0191377

Hong, Y. K., Mcmasters, K. M., Egger, M. E., & Ajkay, N. (2018). Ductal carcinoma in situ: Current trends, controversies, and review of literature. The American Journal of Surgery, 216(5), 998–1003. 10.1016/j.amjsurg.2018.06.013

Jia, L., Yang, T., Gu, X., Zhao, W., Tang, Q., Wang, X., Zhu, J., & Feng, Z. (2019). Translation elongation factor eEF1Bα is identified as a novel prognostic marker of gastric cancer. International Journal of Biological Macromolecules, 126, 345–351. 10.1016/j.ijbiomac.2018.12.126

Kim, J., Chubatsu, L. S., Admon, A., Stahl, J., Fellous, R., & Linn, S. (1995). Implication of mammalian ribosomal protein S3 in the processing of DNA damage. Journal of Biological Chemistry, 270(23), 13620–13629. 10.1074/jbc.270.23.13620

Lambert, M. W. (2018). Spectrin and its interacting partners in nuclear structure and function. Experimental Biology and Medicine, 243(6), 507–524. 10.1177/1535370218763563

Lambert, M. W. (2019). The functional importance of lamins, actin, myosin, spectrin and the LINC complex in DNA repair. Experimental Biology and Medicine, 244(15), 1382–1406. 10.1177/1535370219876651

Latimer, J. J. (2000). Epithelial Cell Cultures Useful For in Vitro Testing (medium) (Patent 6,074,874).

Latimer, J. J. (2002). Epithelial Cell Cultures Useful For in Vitro Testing (protocol) (Patent 6,383,805).

Latimer, J. J., Johnson, J. M., Kelly, C. M., Miles, T. D., & Beaudry-rodgers, K. A. (2010). Nucleotide excision repair deficiency is intrinsic in sporadic stage I breast cancer. PNAS, 107(50), 21725–21730. 10.1073/pnas.0914772107

Lee, I. H., Sohn, M., Lim, H. J., Yoon, S., Oh, H., Shin, S., Shin, J. H., Oh, S.-H., Kim, J., Lee, D. K., Noh, D. Y., Bae, D. S., Seong, J. K., & Bae, Y. S. (2014). Ahnak functions as a tumor suppressor via modulation of TGFβ/Smad signaling pathway. Oncogene, 33(38), 4675–4684. 10.1038/onc.2014.69

Liu, F., Li, Y., Yu, Y., Fu, S., & Li, P. (2007). Cloning of novel tumor metastasis-related genes from the highly metastatic human lung adenocarcinoma cell line Anip973. Journal of Genetics and Genomics, 34(3), 189–195. 10.1016/S1673-8527(07)60020-4

Maxwell, A. J., Clements, K., Hilton, B., Dodwell, D. J., Evans, A., Kearins, O., Pinder, S. E., Thomas, J., Wallis, M. G., & Thompson, A. M. (2018). Risk factors for the development of invasive cancer in unresected ductal carcinoma in situ. European Journal of Surgical Oncology, 44, 429–435. 10.1016/j.ejso.2017.12.007

Narla, A., & Ebert, B. L. (2010). Ribosomopathies: Human disorders of ribosome dysfunction. Blood, 115(16), 3196–3205. 10.1182/blood-2009-10-178129

Ono, K., Sogawa, C., Kawai, H., Tran, M. T., Taha, E. A., Lu, Y., Oo, M. W., Okusha, Y., Okamura, H., Ibaragi, S., Takigawa, M., Kozaki, K.-I., Nagatsuka, H., Sasaki, A., Okamoto, K., Calderwood, S. K., & Eguchi, T. (2020). Triple knockdown of CDC37, HSP90-alpha and HSP90-beta diminishes extracellular vesicles-driven malignancy events and macrophage M2 polarization in oral cancer. Journal of Extracellular Vesicles, 9(1), 1769373. 10.1080/20013078.2020.1769373

Osugi, Y., Fumoto, K., & Kikuchi, A. (2019). CKAP4 regulates cell migration via the interaction with and recycling of integrin. Molecular and Cellular Biology, 39(16), e00073–19. 10.1128/MCB.00073-19

Park, J. W., Kim, I. Y., Choi, J. W., Lim, H. J., Shin, J. H., Kim, Y. N., Lee, S. H., Son, Y., Sohn, M., Woo, J. K., Jeong, J. H., Lee, C., Bae, Y. S., & Seong, J. K. (2018). AHNAK loss in mice promotes type II pneumocyte hyperplasia and lung tumor development. Molecular Cancer Research, 16(8), 1287 LP – 1298. 10.1158/1541-7786.MCR-17-0726

Partek Inc. (2022). Partek Inc. (2020). Partek® Flow® (Version 10.0) [Computer software]. https://www.partek.com/partek-flow/.

Penzo, M., Montanaro, L., Treré, D., & Derenzini, M. (2019). The ribosome biogenesis—cancer connection. Cells, 8(1), 55. 10.3390/cells8010055

Qi, M., Zhang, J., Zeng, W., & Chen, X. (2014). DNAJB1 stabilizes MDM2 and contributes to cancer cell proliferation in a p53-dependent manner. Biochimica et Biophysica Acta (BBA) - Gene Regulatory Mechanisms, 1839(1), 62–69. 10.1016/j.bbagrm.2013.12.003

Ross, P. L., Huang, Y. N., Marchese, J.N., Williamson, B., Parker, K., Hattan, S., Khainovski, N., Pillai, S., Dey, S., Daniels, S., Purkayastha, S., Juhasz, P., Martin, S., Bartlet-Jones, M., He, F., Jacobson, A., & Pappin, D. J. (2004). Multiplexed protein quantitation in Saccharomyces cerevisiae using amine-reactive isobaric tagging reagents. Molecular Cell Proteomics, 3(12), 1154–69. doi: 10.1074/mcp.M400129-MCP200

Roy, R., Huang, Y., Seckl, M. J., & Pardo, O. E. (2017). Emerging roles of hnRNPA1 in modulating malignant transformation. WIREs RNA, 8(6). 10.1002/wrna.1431

Ruggero, D., & Pandolfi, P. P. (2003). Does the ribosome translate cancer? Nature Reviews Cancer, 3(3), 179–192. 10.1038/nrc1015

Sajithlal, G. B., Rothermund, K., Zhang, F., Dabbs, D. J., Latimer, J. J., Grant, S. G., & Prochownik, E. V. (2010). Permanently blocked stem cells derived From breast cancer cell lines. Stem Cells, 28(6), 1008–1018. 10.1002/stem.424

Sanders, M. E., Schuyler, P. A., Simpson, J. F., Page, D. L., & Dupont, W. D. (2015). Continued observation of the natural history of low-grade ductal carcinoma in situ reaffirms proclivity for local recurrence even after more than 30 years of follow-up. Modern Pathology, 28(5), 662–669. 10.1038/modpathol.2014.141

Siegel, R. L., Miller, K. D., Wagle, N. S., & Jemal, A. (2023). Cancer statistics, 2023. CA: A Cancer Journal for Clinicians, 73(1), 17–48. 10.3322/caac.21763

Sjakste, N., Sjakste, T., & Vikmanis, U. (2004). Role of the nuclear matrix proteins in malignant transformation and cancer diagnosis. Experimental Oncology, 26(3), 170–178.

Smith, J. R., Clarke, P. A., de Billy, E., & Workman, P. (2009). Silencing the cochaperone CDC37 destabilizes kinase clients and sensitizes cancer cells to HSP90 inhibitors. Oncogene, 28(2), 157–169. 10.1038/onc.2008.380

Spencer, V. A., Samuel, S. K., & Davie, J. R. (2001). Altered profiles in nuclear matrix proteins associated with DNA in situ during progression of breast cancer cells. Cancer Research, 61(4), 1362–1366.

Treweek, T. M., Meehan, S., Ecroyd, H., & Carver, J. A. (2015). Small heat-shock proteins: important players in regulating cellular proteostasis. Cellular and Molecular Life Sciences, 72(3), 429–451. 10.1007/s00018-014-1754-5

Vaarala, M. H., Porvari, K. S., Kyllönen, A. P., Mustonen, M. V. J., Lukkarinen, O., & Vihko, P. T. (1998). Several genes encoding ribosomal proteins are over-expressed in prostate-cancer cell lines: Confirmation of L7a and L37 over-expression in prostate-cancer tissue samples. International Journal of Cancer, 78(1), 27–32. 10.1002/(SICI)1097-0215(19980925)78:1<27::AID-IJC6>3.0.CO;2-Z

van Seijen, M., Lips, E. H., Thompson, A. M., Nik-Zainal, S., Futreal, A., Hwang, E. S., Verschuur, E., Lane, J., Jonkers, J., Rea, D. W., & Wesseling, J. (2019). Ductal carcinoma in situ: to treat or not to treat, that is the question. British Journal of Cancer, 121(4), 285–292. 10.1038/s41416-019-0478-6

Visus, C., Ito, D., Dhir, R., Szczepanski M. J., Chang, Y. J., Latimer, J. J., Grant, S. G., & DeLeo, A.B. (2011). Identification of hydroxysteroid (17β) dehydrogenase type 12 (HSD17B12) as a CD8^+^ T-cell-defined human tumor antigen of human carcinomas. Cancer Immunology Immunotherapy, 60(7), 919–929. 10.1007/s00262-011-1001-y

Wang, S., & Zhang, Y. (2020). HMGB1 in inflammation and cancer. Journal of Hematology & Oncology, 13(1), 116. 10.1186/s13045-020-00950-x

Wang, W., Nag, S., Zhang, X., Wang, M.-H., Wang, H., Zhou, J., & Zhang, R. (2015). Ribosomal proteins and human diseases: Pathogenesis, molecular mechanisms, and therapeutic implications. Medicinal Research Reviews, 35(2), 225–285. 10.1002/med.21327

Wend, P., Runke, S., Wend, K., Anchondo, B., Yesayan, M., Jardon, M., Hardie, N., Loddenkemper, C., Ulasov, I., Lesniak, M. S., Wolsky, R., Bentolila, L. A., Grant, S. G., Elashoff, D., Lehr, S., Latimer, J. J., Bose, S., Sattar, H., Krum, S. A., & Miranda-Carboni, G. A. (2013). WNT10B/β-catenin signalling induces HMGA2 and proliferation in metastatic triple-negative breast cancer. EMBO Molecular Medicine, 5(2), 264–279. 10.1002/emmm.201201320

Xiong, J., Li, Y., Tan, X., & Fu, L. (2020). Small heat shock proteins in cancers: Functions and therapeutic potential for cancer therapy. International Journal of Molecular Sciences, 21(18), 6611. 10.3390/ijms21186611

Yang, Z.-Y., Jiang, H., Qu, Y., Wei, M., Yan, M., Zhu, Z.-G., Liu, B.-Y., Chen, G.-Q., Wu, Y.-L., & Gu, Q.-L. (2013). Metallopanstimulin-1 regulates invasion and migration of gastric cancer cells partially through integrin β4. Carcinogenesis, 34(12), 2851–2860. 10.1093/carcin/bgt226

Yao, Y., Dong, Y., Lin, F., Zhao, H., Shen, Z., Chen, P., Sun, Y.-J., Tang, L.-N., & Zheng, S.-E. (2009). The expression of CRM1 is associated with prognosis in human osteosarcoma. Oncol Rep, 21(1), 229–235. 10.3892/or_00000213

Yuan, S., Liu, Z., Xu, Z., Liu, J., & Zhang, J. (2020). High mobility group box 1 (HMGB1): a pivotal regulator of hematopoietic malignancies. Journal of Hematology & Oncology, 13(1), 91. 10.1186/s13045-020-00920-3

Zhan, Y., Melian, N. Y., Pantoja, M., Haines, N., Ruohola-Baker, H., Bourque, C. W., Rao, Y., & Carbonetto, S. (2010). Dystroglycan and mitochondrial ribosomal protein L34 regulate differentiation in the Drosophila eye. PLOS ONE, 5(5), e10488.

Zhao, Z., Xiao, S., Yuan, X., Yuan, J., Zhang, C., Li, H., Su, J., Wang, X., & Liu, Q. (2017). AHNAK as a prognosis factor suppresses the tumor progression in glioma. Journal of Cancer, 8(15), 2924–2932. 10.7150/jca.20277

Zink, D., Fischer, A. H., & Nickerson, J. A. (2004). Nuclear structure in cancer cells. Nature Reviews Cancer, 4(9), 677–687. 10.1038/nrc1430

