## Supplemental Data for "Cancer-Specific Alterations in Nuclear Matrix Proteins Determined by Multi-omics Analyses of Ductal Carcinoma *in Situ*"

**Extended View Table 1. Clinical characteristics of non-diseased and breast cancer explants used in this study.**

| Cell Culture | Tumor Stage |
|---|---|
| JL-BRL-6 (Breast Reduction mammoplasty culture) | |
| JL-Contra-3 Contralateral Breast culture for DCIS-3 | |
| JL-DCIS-3 | Stage 0 |
| JL-BTL-4 | Stage I |
| JL-BTL-8 | Stage I |
| JL-BTL 33 | Stage I |
| JL-BTL-37 | Stage I |
| JL-BTL-9 | Stage II |
| JL-BTL-10 | Stage II |
| JL-BTL-29 | Stage II |
| JL-BTL-46 | Stage II |
| JL-BTL-12 | Stage III |
| JL-BTL-21 | Stage IV |
| JL-BTL-60 | Stage IV |
| MDA-MB231 | Stage IV |
| MCF-7 | Stage IV |

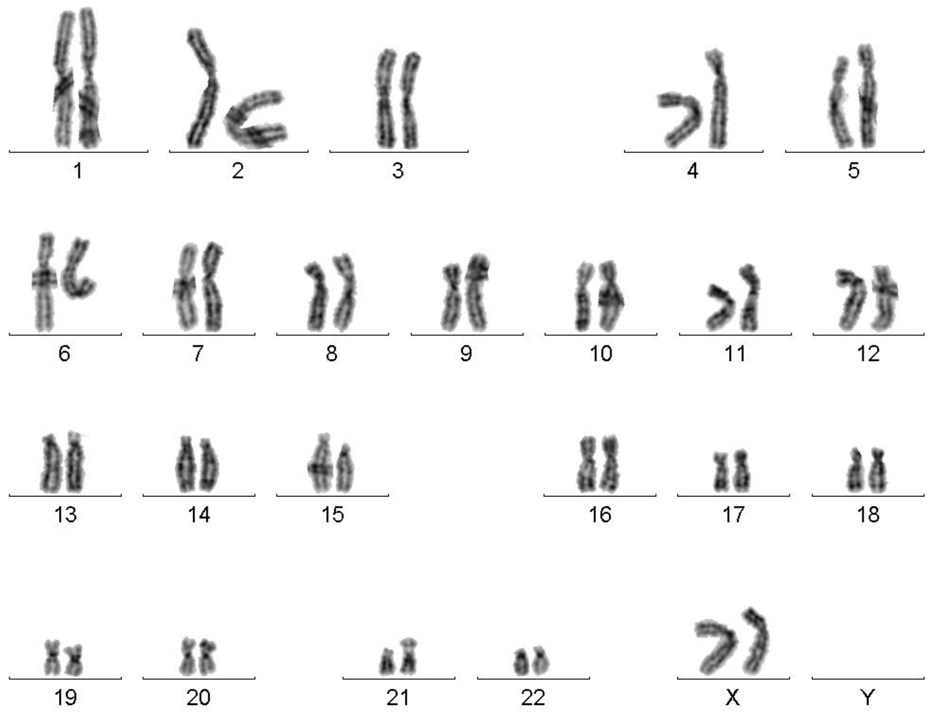


**Fig EV1 A.** Karyotype of JL-Contra-3 cell line showing a diploid normal female chromosome complement

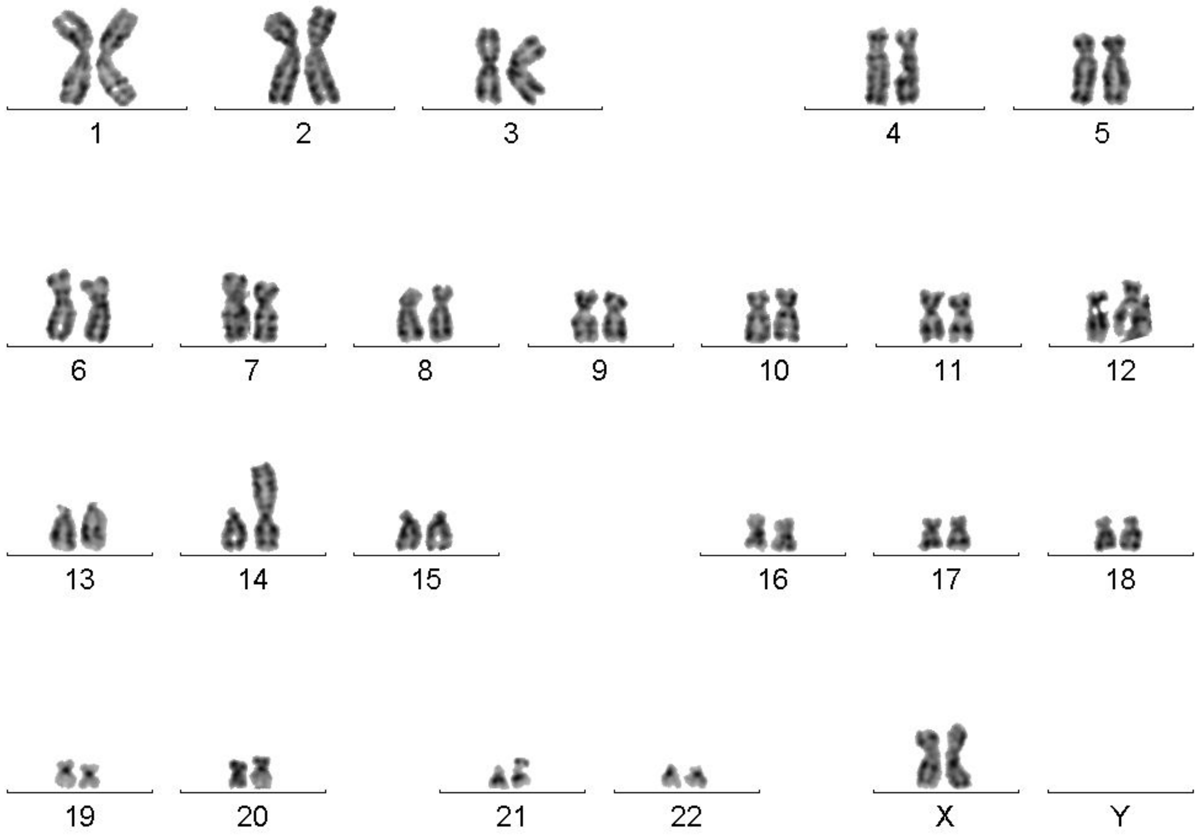


**Fig EV1 B.** Karyotype of JL-DCIS-3 showing a derivative chromosome 14 with an extra copy of 5q. This karyotype represents a prominent subclone of this patient-derived culture.

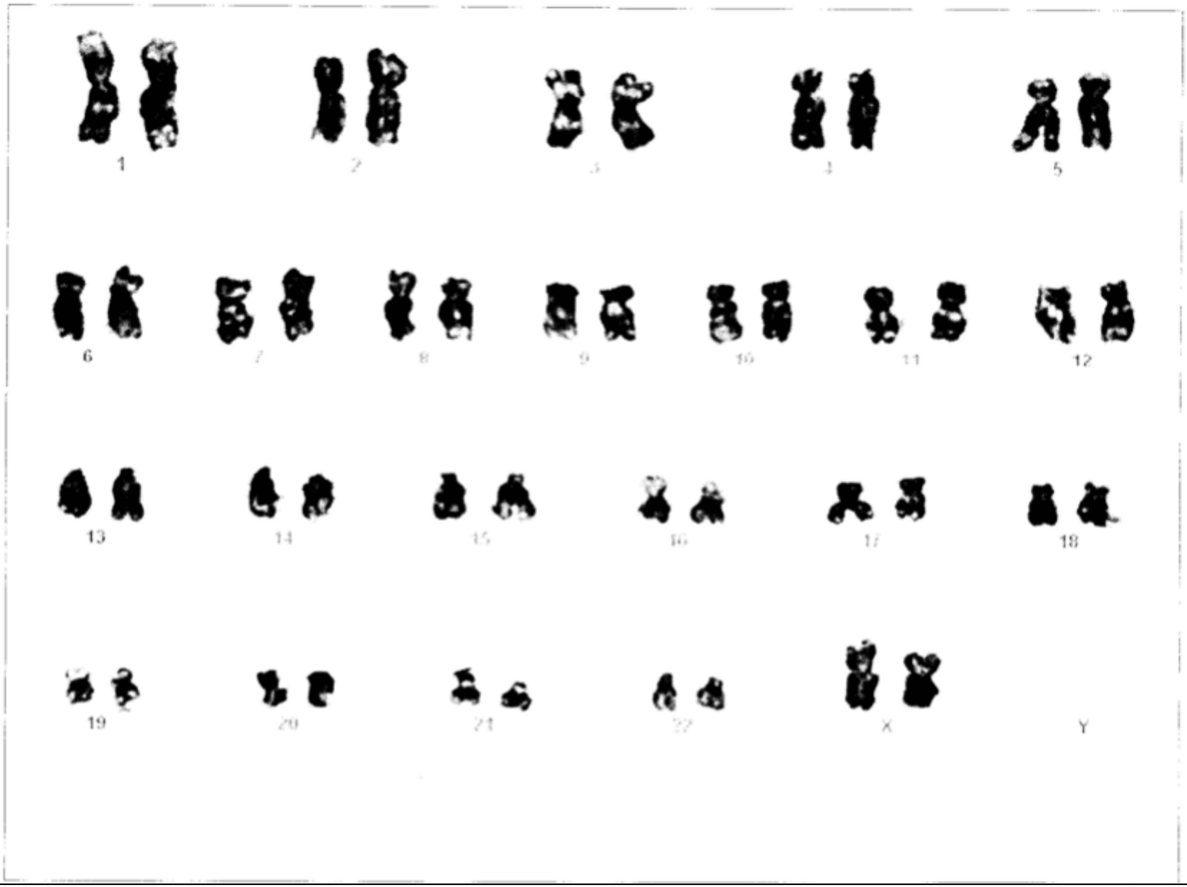


**Fig EV1 C.** Karyotype of JL-BRL-6 showing a diploid normal female chromosome complement.
